# Temporal phenomic predictions from unoccupied aerial systems can outperform genomic predictions

**DOI:** 10.1101/2021.10.06.463310

**Authors:** Alper Adak, Seth C. Murray, Steven L. Anderson

**Affiliations:** Dept. of Soil and Crop Sciences, Texas A&M Univ., College Station, TX77843-2474, USA; Dept. of Environmental Hort., Institute of Food and Agricultural Sciences, Mid-Florida Research and Education Center, University of Florida, Apopka, FL, USA

**Keywords:** high throughput phenotyping, phenomic prediction, genomic prediction, plant breeding

## Abstract

A major challenge of genetic improvement and selection is to accurately predict individuals with the highest fitness in a population without direct measurement. Over the last decade genomic predictions (GP) based on genome-wide markers have become reliable and routine. Now phenotyping technologies, including unoccupied aerial systems (UAS also known as drones), can characterize individuals with a data depth comparable to genomics when used throughout growth. This study, for the first time, demonstrated that the prediction power of temporal UAS phenomic data can achieve or exceed that of genomic data. UAS data containing red-green-blue (RGB) bands over fifteen growth time points and multispectral (RGB, red-edge and near infrared) bands over twelve time points were compared across 280 unique maize hybrids. Through cross validation of untested genotypes in tested environments (CV2), temporal phenomic prediction (TPP) outperformed GP (0.80 vs 0.71); TPP and GP performed similarly in three other cross validation scenarios. Genome wide association mapping using area under temporal curves of vegetation indices (VIs) revealed 24.5 percent of a total of 241 discovered loci (59 loci) had associations with multiple VIs, explaining up to 51 percent of grain yield variation, less than GP and TPP predicted. This suggests TPP, like GP, integrates small effect loci well improving plant fitness predictions. More importantly, temporal phenomic prediction appeared to work successfully on unrelated individuals unlike genomic prediction.

## Introduction

To improve genetic gain, plant breeders must phenotype more plants repeatedly during growth allowing higher selection intensity, accuracy, and increased statistical power (1-3). High quality and quantity phenomic data is essential to develop widely applicable prediction models (e.g., phenomic predictions) to predict yield across growing environments and conditions in the near future (4). To date, few phenomic data sets, approaches and applications have been reported, especially those applied in a breeding context.

Organismal fitness, such as terminal grain yield in crops, is a cumulative response of genetics (G), the environment (E), management (M) and integrated GxExM interactions temporally throughout growth. To predict cumulative fitness of an individual organism without direct measurement of that individual’s fitness, proxies such as genetic markers are used, to link measurements of relatives and predict fitness with breeding values. Traditional best linear unbiased prediction (BLUP) derived breeding values (5) were modified by (6) where genotypic marker data of parental inbreds was combined with the yield data of the related single cross hybrids to predict yield performance of the single cross hybrids, known as genomic BLUP (GBLUP). However, prediction accuracies dropped dramatically when yield of unknown (previously untested) parental lines derived hybrids was predicted (7, 8). Various genomic based statistical models have been developed after the traditional GBLUP approach with advent of genomic technology (9-11). These methods have been applied extensively as genome wide marker facilitated selection also known as genomic selection in plants (12). Predicting the performance of previously untested genotypes in both tested and untested environments remains the central problem in plant breeding selections, and new approaches to addressing this challenge are needed. Genomic selection to estimate genotype fitness, as measured by terminal grain yield, relies on manually collected phenotype data which is resource intensive to collect. Phenotypic characteristics of cumulative complex traits are often not accurately predicted in GS because of (i) the different interplays of genes on phenotype throughout different growth stages, (ii) different effect sizes of the same genetic markers on phenotype of complex traits at different growth stages and (iii) different sources of phenotypic variation of the complex traits at different growth stages (13-21). Tools that can inexpensively evaluate individuals throughout growth, as they interact with their environment, would therefore be a valuable addition to predicting an organism’s fitness. Unoccupied aerial systems (UAS) are now able to provide these insights, frequently evaluating individuals temporally throughout growth. However, to date, fitness predictions from UAS alone have not been compared to the standard method of genomic prediction.

To evaluate fitness prediction of UAS based phenomics tools, the breeding value of each hybrid must be produced, these can be estimated from temporal VIs and structural measurements (canopy height) collected temporally throughout growth. Correlations between temporal VIs with yield and flowering times, as well as machine learning models can investigate predictive abilities for fitness traits (yield and flowering times). Phenomic predictions made from temporal vegetation indices and canopy height can be compared with traditional genomic predictions. Ultimately, major causal loci underlying phenomic predictions success for complex traits can be useful to understand underlying biology of organismal fitness over growth. Here we report phenomic data-driven selection for complex traits in maize breeding. We conducted UAS surveys with multispectral and RGB sensors to collect image-based temporal predictors throughout maize growth stages. We compared phenomic based prediction accuracy to that of genomic prediction, explored temporal shifts in image-based phenotypic variation explained by genome wide markers, and conducted association mapping utilizing temporal image-based phenotypes to identify biologically important loci.

## Materials and Methods

Using the Genome to Fields initiative’s 2017 germplasm, 280 unique maize hybrids were grown under optimal management (OM) and 230 were grown under stressed management (SM, no irrigation, low fertilizer) near College Station, Texas. Two replications were used in a randomized complete block design with each hybrid grown as two consecutive row plots.

### UAS surveys and image processing

A Phantom 3 Professional rotary-wing UAS, equipped with a 12-megapixel red-green-blue (RGB) DJI FC300X camera, flown 25 meters above the ground (TPP_RGB). Additionally, a Tuffwing UAS equipped with a MicaSense RedEdge-MX multispectral camera was flown 120 meters above the ground (TPP_Multi). Images were collected with 80% forward and side overlap in both surveys. Raw images were processed in Agisoft Metaphase Professional software (https://www.agisoft.com/) to generate the 3D point clouds and orthomosaics (***SI appendix*, Table S1)** (22).

### Phenomic data extraction pipeline

Environmental Systems Research Institute, Inc. (ESRI) shape file were constructed using R/UAStools::plotshpcreate function (23) and applied to each survey’s respective orthomosaic (.tif files) and 3D point clouds (.las or .laz files) to extract plot level image based phenotypes. Vegetation indices (***SI appendix*, Table S2)** for each flight date were extracted using the *FIELDImageR* package (24) for each UAS survey (***SI appendix, SI Materials and Methods*)**. Plot based 99^th^ percentile temporal plant heights (canopy height measurement; CHM) were extracted from 3D point clouds following the methods of (16) (***SI appendix, SI Materials and Methods*)**.

### Experimental design and nested model for phenomic data

To analyze the temporal VIs and CHM, a custom nested design was applied to raw data of each VI and CHM belonging to each row plot in OM and SM, where experimental design and maize hybrids were treated as nested within drone times (***SI appendix, SI Materials and Methods*)**. Hybrids nested within pedigree results were used to predict GY, DTA, DTS, and PHT within and between the trials.

### Machine learning based phenomic prediction models

Manually collect phenotypes (GY, DTA, DTS and PHT) were predicted using linear, elastic net, ridge, lasso, and random forest regressions using the TPP_RGB and TPP_Multi image-based phenotypes. Prediction models were trained using a random sampling of 70% of the common maize hybrids (tested genotypes) across the two management environments. The remaining 30% were used as the validation dataset (untested genotypes). Models were trained using OM trial (tested environment) while the SM trial served as the untested environment. Four cross validation schemes (CVs) were conducted as follows: (i) tested genotypes in tested environment (CV1), (ii) untested genotypes in tested environment (CV2), (iii) tested genotypes in untested environment (CV3), and (iv) untested genotypes in untested environment (CV4) (25). Phenomic prediction models and prediction steps are available in the ***SI appendix, SI Materials and Methods***.

### Association mapping for phenomic data

The image-based vegetation indices and Weibull_CHM were converted to cumulative area under curve (AUC) values and used as trait data in a genome wide association study (GWAS) **(*SI appendix, SI Materials and Methods*)**. Association mapping was conducted using 158 maize hybrids and 101,100 genotyping by sequencing (GBS) SNP markers, implementing three multiple loci test methods; (i) fixed and random model circulating probability unification (FarmCPU) (26), (ii) multiple loci mixed model (MLMM) (27), and (iii) bayesian-information and linkage-disequilibrium iteratively nested keyway (BLINK) (28) (***SI appendix, SI Materials and Methods*)**. Linkage disequilibrium (LD) estimates were used to identify candidate genes within LD blocks (*R*^2^ ≥ 0.8) of colocalized SNPs (***SI appendix*, Fig. S1**).

### Genomic prediction for phenomic data

Genome-wide prediction was applied to 540 image-based phenotypes (35 VIs and CHM belonging to 16 flight times) of the 158 maize hybrids in TPP_RGB using 153,252 SNPs, temporal genomic prediction model was explained in ***SI appendix, SI Materials and Methods***.

### Phenomic prediction versus genomic prediction

GBS marker data (GP) and two sets of phenomic data (TPP_RGB and TPP_Multi) were used to conduct genomic prediction and phenomic prediction for maize grain yield (GY). A total of 118 G2F maize hybrids were used to compare the predictive ability between the genomic and phenomic data sets. Four cross validation schemes were applied as explained in “*Machine learning based phenomic prediction models”* section. Additional details regarding phenomic prediction versus genomic prediction are available within the ***SI appendix, SI Materials and Methods***.

## Results

### Variance decomposition and repeatability estimates demonstrate UAS sensor-based phenotypes were genetically stable

Variance component decomposition of the 83 sensor-based VIs (35 RGB and 54 multispectral) demonstrated UAS sensor-based data was statistically repeatable and biologically meaningful with a genetic basis. The rotary-wing equipped with an RGB (3 band, 12 MP) sensor flown at 25 m resulted in ∼1 cm pix^-1^ image resolution and had higher repeatability than the Tuffwing platform equipped with a multispectral (5 band, 3.8 MP) sensor flown at 120 m (∼8 cm pix^-1^). The main source of phenotypic variation for both platforms was explained by the temporal flight component (*β*_*i*_ component in *Eq*. 1) of the nested design (31-96%) showing a temporal plasticity of maize spectral reflectance signatures throughout the plants growth cycle (***SI appendix*, Figs. S2 and S3**). Genetic variance (*Ω*_*i*(*j*)_ component in *Eq*. 1) was slightly greater for the higher resolution-low altitude RGB (1.5 - 5.2%; TR: 0.46 - 0.77) phenotypes compared to the lower resolution-high altitude RGB (1.1 - 4.5%; TR: 0.26 - 0.66) and lower resolution-high altitude multispectral (0.5 - 3.4%; TR: 0.28 - 0.62) phenotypes (***SI appendix*, Figs. S2 and S3**). The repeatability estimates over the 35 RBG phenotypes were highly correlated (r=0.71) between the two sensor systems, although repeatability was improved by 0.08 on average, when implementing the higher resolution-low altitude RGB platform. Noticeable improvements in repeatability estimates (>0.1) were achieved for 13 RGB VIs and 6 VIs repeatability were reduced (<0.06) when implementing the higher resolution-low altitude RGB platform (***SI appendix*, Figs. S2 and S3**). Overall, significant genetic variation was attributed to all VIs on both platforms, useful in predictive modeling of important agronomic traits such as grain yield, flowering times, and plant height (***SI appendix*, Fig. S4**).

### Temporal correlation

Temporal correlation between UAS survey dates of the VIs derived from the higher-resolution-low altitude RGB demonstrated that 14 of 35 RGB-derived VIs achieved a correlation above 0.50 (up to 0.61) to GY (**SI appendix, Fig. S5**). However, the 14 RGB- and 40 multispectral-derived VIs from the lower resolution-high altitude multispectral achieved correlations above 0.50 (up to 0.70) to GY (**SI appendix, Fig. S6**). Sensor-based VIs correlations with GY varied depending on the flight times. High correlations were found for VIs belonging to certain time points in both TP_RGB and TPP_Multi demonstrating that temporal VIs tend to synchronize with GY in maize hybrids indicating potential source for predicting yield.

### Phenomic prediction using high dimensional UAS data

Temporal breeding values of each pedigree at each timepoint in TPP_RGB and TPP_Multi followed unique trajectories (***SI appendix*, Figs. S7 and S8)** visually discriminating low, mid, and high yielding maize hybrids. Phenotype data of VIs at different time points had different discriminative ability for yield. This led us to test the predictive ability of two phenomic data derived from different sensors and resolutions utilizing the different prediction models.

The three machine learning models improved prediction accuracy (>90%) for all four agronomic traits (GY, DTA, DTS, and PHT) compared to the linear model when temporal phenotypes in TPP_RGB and TPP_Multi phenomic data were used as predictors (**Fig. 1**). The linear models had the highest prediction errors (RMSE and MAE) and lowest R^2^ (***SI appendix*, Fig. S9**). Ridge regression was the highest performing model for predicting GY regardless of the phenomic data sets; resulting in the best prediction performances for untested genotypes in tested environment (CV2), tested genotypes in untested environment (CV3), and untested genotypes in untested environment (CV4) (**Fig. 1**). Ridge regression best predicted GY for CV2 using the low-resolution multispectral sensor (TPP_Multi), while ridge regression also best predicted GY for tested and untested genotypes in untested environment cross validations (CV3 and CV4) using the high resolution RGB sensor (TPP_RGB; **Fig. 1**). Furthermore, ridge regression achieved the greatest prediction accuracy for the flowering times and plant heights utilizing the high resolution RGB UAS (**Fig. 1**). Prediction accuracies were higher in the most challenging CVs (CV3 and CV4) when TPP_RGB was used to predict GY, DTA, DTS, and PHT. These results demonstrate that the reduction in resolution, increased spectral bands, and increased sensor cost of incorporating the multispectral bands did not significantly improve model performance.

**Fig. 1.**
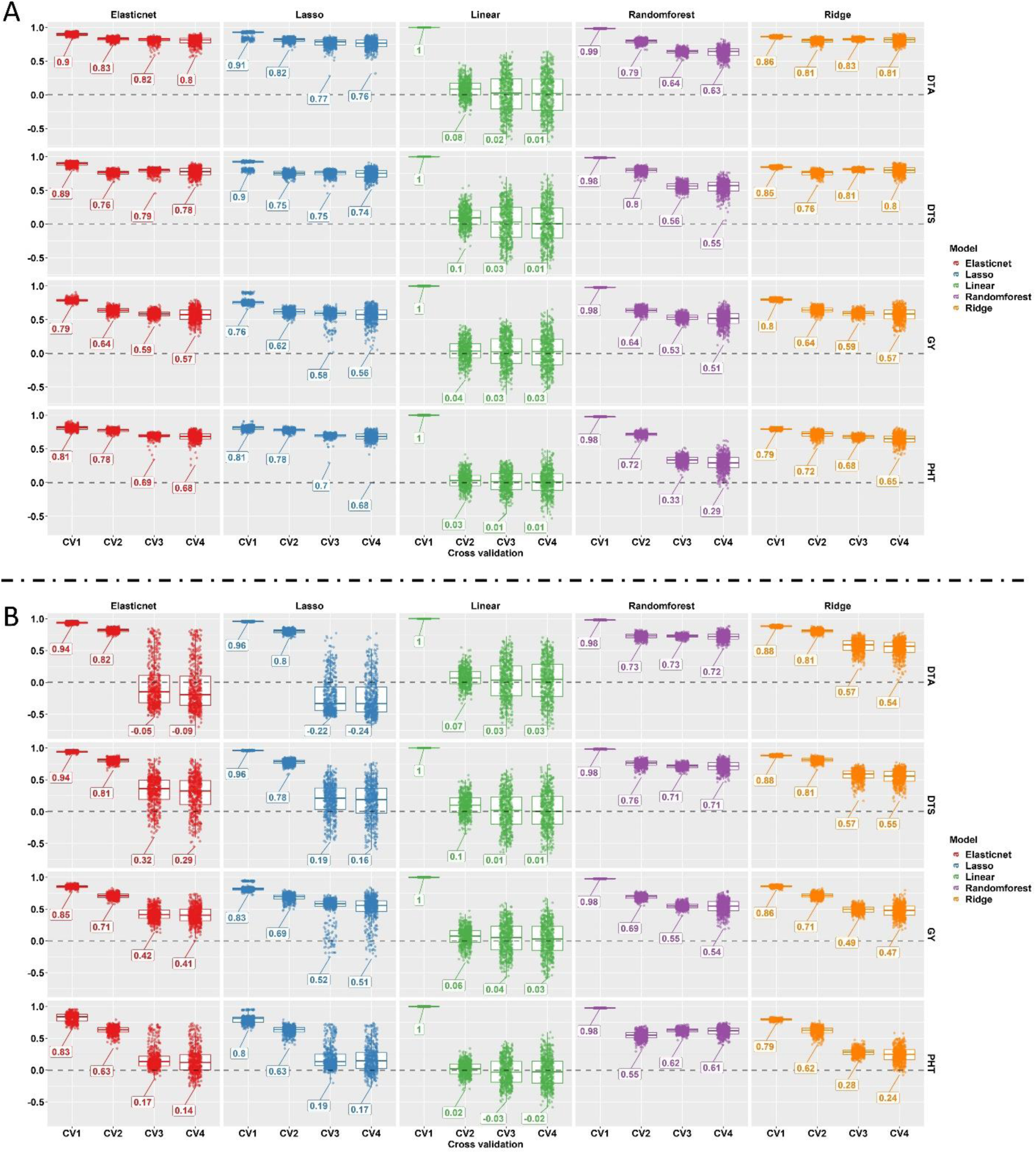
Shows the prediction accuracy (on the y axis) of the phenomic prediction obtained by each model for four cross validation schemes (on the x axis) belonging to each predicted variable (from left to right) in phenomic prediction: (A) the prediction performance of TPP_RGB phenomic data derived from HTP platform including 25-meters elevation with RGB sensor. (B) the prediction performance of TPP_Multi phenomic data derived from HTP platform including 120-meters elevation with multispectral sensor. Ridge regression performed best overall for the most challenging cross-validation schemes, CV3 and CV4, compared to other prediction models when TPP_RGB was used. Whereas, Randomforest performed best overall for the most challenging cross-validation schemes, CV3 and CV4, compared to other prediction models when TPP_Multi was used.

### Variable importance scores of the machine learning models

To understand potential biological causes behind the most accurate predictions, variable importance scores were derived from the prediction models to identify critical predictor/time point combinations for TPP_RGB and TPP_Multi phenomic data sets (***SI appendix*, Figs. S10 and S11)**. Different contributions of VIs and Weibull_CHM at multiple time points were important among both phenomic datasets in the prediction of GY, DTA, DTS and PHT (***SI appendix*, Figs. S10 and S11)**. For instance, the TPP_RGB red chromatic coordinate index (RCC) and TPP_Multi modified nonlinear index values (MNLI) belonging to various time points, either before or after flowering times, for all predicted variables were identified by all machine learning models consistently and are therefore critical VI/timepoints combinations for all predicted variables (***SI appendix*, Figs. S10 and S11)**. This demonstrates the ability of machine learning models to identify important image-based phenotypes for future UAS surveying efforts and provides foundational insight towards understanding the biological importance of images-based phenotypes within a plant’s growth cycle.

### Genome wide association mapping results

To gain further insight into biological significance of successful predictions, GWAS peaks were identified using area under curve values (***SI appendix*, Fig. S12)** of each high resolution VI and Weibull_CHM in the TPP_RGB phenomic data set (***SI appendix*, Fig. S13)**. A total of 241 GWAS peaks were identified across the 36-temporal image-based phenotypes in TPP_RGB. Five genomic regions had significant loci for VIs and candidate genes of relevant interest **(*SI appendix, SI Results*)**. Two genomic regions were identified as hotspots (fourth bin in chr2 and eighth bin in chr4) having GWAS peaks belonging to 24 VIs discovered across the three GWAS models (***SI appendix*, Fig. S14 and Dataset S1**). A 15 kb genomic distance around the GWAS peaks was scanned to determine candidate genes based on the calculated LD decay (***SI appendix*, Fig. S1**). LD patterns of both hotspots were visualized along with six candidate genes with functions described in ***SI appendix*, Fig. S14**.

A hotspot was identified at 36,828,844 bp on chromosome 2 (*chr2_1*), identified by the excessive red, modified green red, normalized difference, Normalized green red difference, and visible atmospherically resistant indices by the three GWAS models consistently explaining 8-13% phenotypic variation (**Dataset S1**). The *chr2_1* peak is inside *GRMZM2G023204* (chr2:36827859..36,829,876; B73 RefGen_v4), a putative protein kinase domain that catalyzes the function of protein kinases. Another candidate gene (∼4kb away from *chr2_1*) is *GRMZM2G021560* (*pebp25;* chr2:36,779,809..36,782,444; B73 RefGen_v4) a member of phosphatidylethanolamine-binding proteins (PEBPs) that regulate floral transitions (29) as well as that *GRMZM2G021560* found to be expressed at the early vegetative stage (eg. third leaf stage) (30). Integrating GWAS with temporal phenotypes (TPP RGB), loci controlling the temporal VIs explained the phenotypic variations of multiple VIs revealing the pleiotropic effects of the loci. Additional candidate genes for other hotspots are discussed in ***SI appendix***.

### Genomic prediction results of temporal phenomic data

Genomic prediction results of temporal VI’s identified specific time points for each of the high-resolution VIs in TPP_RGB had varying ability to be predicted in cross validation (**Fig. 2**). Prediction accuracy showed flowering was the most (and in a few cases least) predictable by genomic markers for many VI’s likely because of differential emergence of tassels **(Fig. 2)**. It was surprising that time points prior to flowering in some cases had relatively similar or higher prediction accuracy than those at flowering time **(Fig. 2)**. Overall, sensor-based VIs were predictable at different time points using whole genome markers but estimated different phenotypic effect sizes **(Fig. 2)**. This demonstrates that genetic makers estimated changing effects sizes revealing the plasticity of temporal VIs that are more explanatory to monitor the interactions between genetic background of plants and their growing environments across plant growth.

**Fig. 2.**
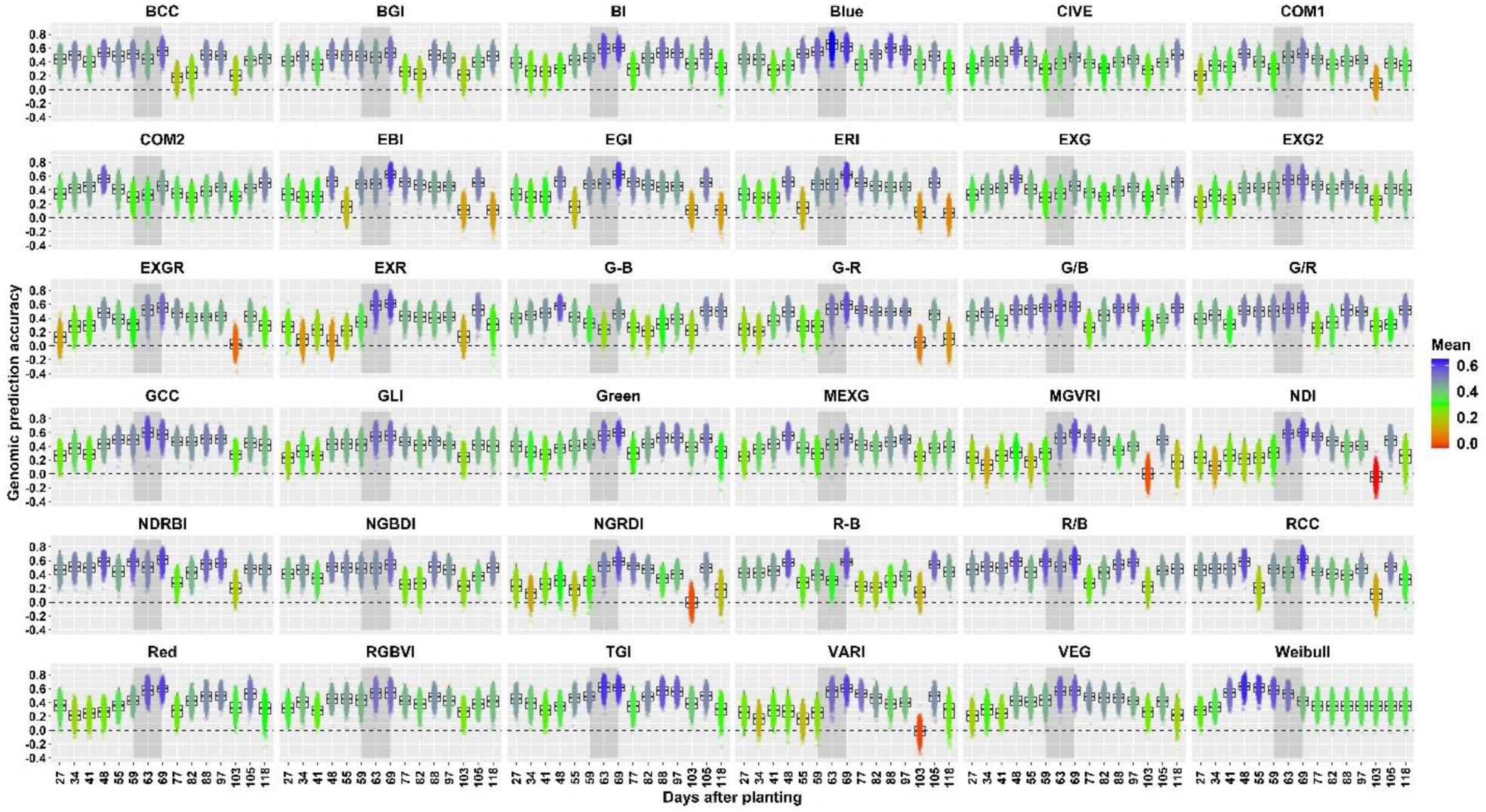
Each box plot shows the genomic prediction accuracy results belonging to each time points of each temporal trait in TPP_RGB, each contains 500-prediction accuracies. Y axis shows the prediction accuracy and x axis shows the flight date as days after planting time. Each box plot was colored based on their mean. Heatmap color scale was given in the figure legend changing between 0 to 0.6. Gray shading in each represents flowering time. Different time points of temporal traits were found to have different response to genetic markers across growth stages of plant development.

### Genomic prediction vs phenomic prediction

Grain yield (GY) prediction ability of phenomic and genomic approaches were compared between both phenomic data sets (TPP_RGB and TPP_Multi) and genomic data (genomic prediction, GP). Comparing model prediction accuracies for untested genotypes in tested environment (CV2), low resolution multispectral (TPP_Multi) outperformed 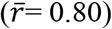 both genomic prediction 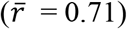 and high resolution RGB 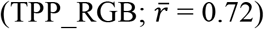 (**Fig. 3**). Comparing model prediction accuracies for untested genotypes in untested environment (CV4), genomic prediction and RGB high resolution phenomic selection supplied similar prediction accuracies 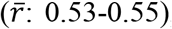, while low resolution with multispectral sensor based HTP supplied a lower prediction accuracy (**Fig. 3**). Overall, the phenomic prediction platforms used in this study were largely able to predict better (CV2), or equivalent to, genomic prediction (CV1 and CV3) depending on which of the four cross validation schemes is evaluated **(Fig. 3)**. However, genomic prediction outperformed phenomic prediction when predicting known genotypes in unknown environments (CV3). Combining both UAS measures (TPP_RGB and TPP_Multi) using ridge regression did not further improve prediction accuracies (data not shown).

**Fig. 3.**
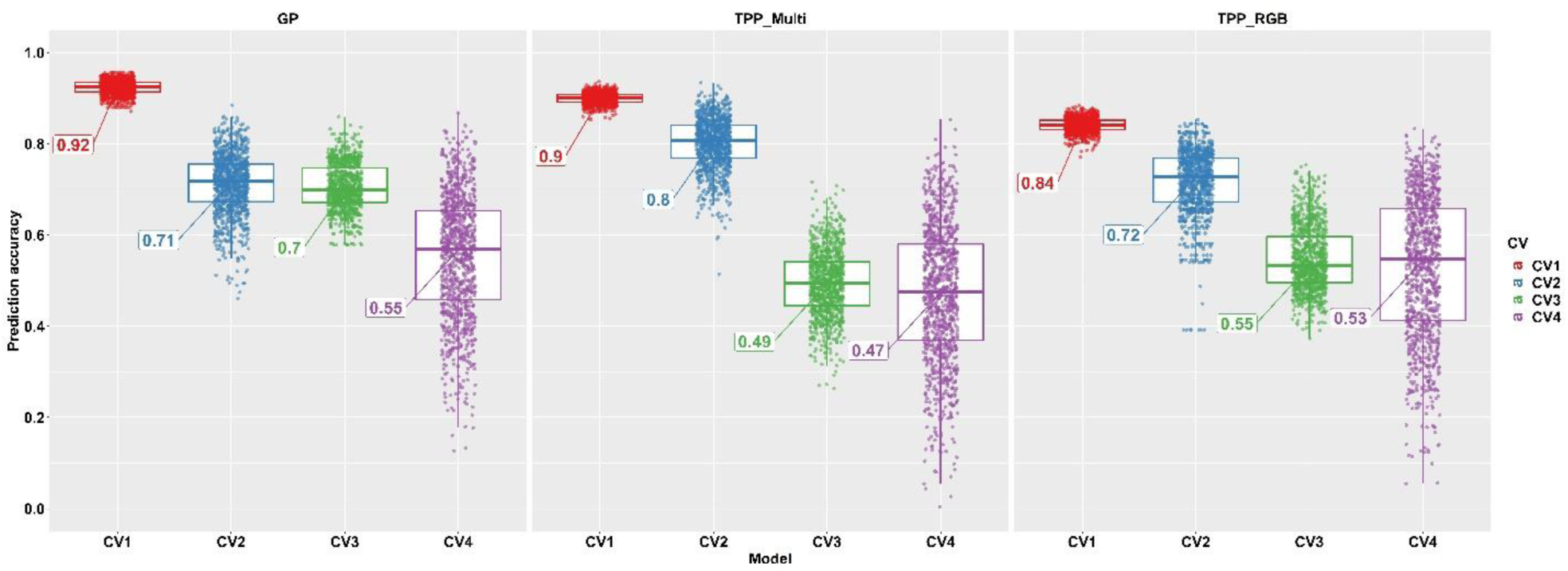
shows the prediction accuracy results of yield belonging to the three models. GP represents the prediction accuracy of genomic prediction, TPP_Multi represents the prediction accuracy of phenomic prediction using the Vis derived from the multispectral images with low resolution, TPP_RGB represents the prediction accuracy of phenomic prediction using the VIs derived from the RGB images with high resolution. Four cross validation schemes were used: predicting tested genotypes in tested environments (CV1), predicting untested genotypes in tested environments (CV2), tested genotypes in untested environments (CV3), and untested genotypes in untested environments (CV4). Phenomic prediction predicted the grain yield (GY) of maize hybrids better in CV2 than genomic prediction. Prediction accuracies were close to each other in CV3 and CV4.

## Discussion

Field-based high-throughput phenotyping technologies, such as drones, are able to provide phenome-wide measurements of plants in much the same way that high-throughput sequencers have provided genome-wide data. Uniquely, phenotyping technologies can screen high numbers of plots repeatedly through the growing period resulting in not only high spatial resolution but also high temporal resolution, helping dissect how different genotypes respond to their environments to maximize fitness in near real-time (***SI appendix*, Figs. S7 and S8)**.

As new temporal phenomic markers are difficult to independently measure and validate, one of the first approaches to evaluate phenomic marker utility is to look at heritability/repeatability values over different replicates and environments. This approach is not needed for genomic markers which do not vary over replicates and environments and theoretically have a repeatability near 1, but are also unable to capture environmental interaction in real time. Temporal repeatability (*Eq. 3*) of VIs were moderate, above ∼0.5 for TPP_RGB (***SI appendix*, Fig. S2)** and between 0.26 and 0.66 for TPP_Multi (***SI appendix*, Fig. S3)**. Temporal repeatability relied on variation across plant development, biologically more meaningful than using genotypic variation which is static at every time point. Temporal variation captured by drones assesses temporal genotypic variation jointly over time via nested design (*Eq. 2*). Previously, repeatability has only been calculated between different vegetation indices/CHM and yield at a single time point (16, 31-37); disregarding the temporal genotypic variation occurring across plant growth. Furthermore, previous studies used either one or a limited number time points and analyzed each time point separately.

High dimensional and temporal resolution phenomic data used in predictive plant breeding integrated with high throughput genotyping data discovered underlying genetic causes for many important temporal VI features. For instance, pleiotropy discovered via GWAS identified specific loci controlling more than one VIs (***SI appendix*, Figs. S13 and S14, Dataset S1)**. In addition, genomic prediction of temporal VI phenotypes proved that estimated effects of each marker varied through time, causing different prediction accuracy results for temporal phenotypes of the same VIs **(Fig. 2)**. Therefore, instead of depending on discrete genome wide markers as predictors for yield, temporal phenotype data formed by estimated temporal marker effects could better predict certain scenarios (e.g., untested genotypes in tested environment). Predicting grain yield of untested genotypes in a tested environment is an important scenario for public breeding programs because lines developed in public breeding programs are mostly targeted for specific environments. So that **figure 3** proved that TPP predicted the grain yield better than GP in CV2 indicating that TPP can be better solution for the public breeding programs for genetic gain. In addition, the predictive ability of TPP in untested genotype untested environments (CV4) was in the same range as that of GS **(Fig. 3**). This is also an important proof of concept that TPP can be used as widely as GP. Genomic prediction methods have been developed over more than a decade and phenomic prediction methods can likewise be improved. Further optimization and improvement of this approach will likely benefit from the integration of novel crop growth models as genomic prediction has (38).

### Phenomic data can predict yield and flowering times via machine learning regressions

Shrinkage factors previously shown as the best performing prediction models when using different hyper parameters have been adapted for predicting both yield (15, 31, 39) and flowering times (15) when different reflection bands were used as predictors. Machine learning models with different regularization parameter settings to predict yield and flowering times (**Fig. 1**) were more accurate than linear-based prediction models (15, 34). This suggests that temporal variation in VIs do not have a linear relationship to predicted variables. This is because linear models tend to overfit when there are increasing numbers of predictors and with fluctuating collinearity between predictors, such as in phenomic data. Linear models are not capable to explain non-linear relationships between predictors and predicted variables.

Tuning regularization parameters of the ridge, lasso and elastic net-based prediction models is a good approach to deal with model overfitting when high dimensional phenomics data are used in prediction. Tuned regularization parameters in ridge, lasso and elastic net models can lessen coefficients, and predict test data more reliably than linear models. For example, pedigree within flight combination (*Ω*_*i*(*j*)_ component in *Eq. 2*) were found to be statistically significant for all VI and CHM (***SI appendix*, Fig. S2)** indicating a temporal interaction among the pedigree across flight times because of fluctuating temporal phenotype values of VIs (***SI appendix*, Figs. S7 and S8**). Nevertheless, a general trend demonstrated that high- and low-yielding pedigrees segregate according to temporal phenotypes of VIs. This reverse correlation of temporal breeding values of the pedigree through time supports the existence of nonlinear relationships, problematic for a linear model to capture. Because of multiple decision tree learning, the random forest model accounts for non-linearity, limiting overfitting.

Phenomic prediction reached up to ∼0.80 for grain yield and flowering time prediction (**Fig. 1**) higher than previously reported prediction accuracies (31-33, 35-37). (31) showed use of raw reflected bands instead of ratios (e.g. vegetation indices) performed better in prediction models. (34) further reported using all bands simultaneously increased prediction accuracy instead of VIs alone. However, reflected bands used in past studies derived from five to nine time points, lower time dimension data than what we generated in our study. This suggests that predictors derived from additional time points could play an important role on increasing the prediction ability of the models; more so than using the predictors as either raw reflectance bands or vegetation indices.

### Genomic prediction for temporal traits can vary depending on the time points of growth

TPP_RGB phenomic data tested using genomic prediction to identify temporal marker effects and their prediction accuracies for each VI and Weibull_CHM throughout time (**Fig. 2**) demonstrated that genomic markers could predict an individual’s VI or Weibull_CHM value through cross validation using other individuals at the same stage. This demonstrated that certain stages and VIs have more genetic determination and are more heritable.

Temporally varying marker effects on the phenotype of VIs resulted in phenotypes at different timepoints of VIs and Weibull_CHM having different correlations with yield (***SI appendix*, Figs. S5 and S6**) as well as different prediction abilities for dependent variables (**Fig. 2**). A dynamic pattern of marker effects as shown here has so far been overlooked in genomic prediction/selection of yield. (4) underlined that predicting the candidate genotype using the phenotype information collected from across multiple environments may be more accurate than using the genetic markers in the prediction model. Similarly, instead of predicting grain yield fitness by whole genome marker effect approaches such as RR-BLUP and GBLUP, including the temporal phenotypic variation occurring across growth into prediction models can result in more accurate fitness prediction as phenomic data already contain temporal marker effects. This study also showed that specific loci can explain different phenotypic variance across more than one derived VI (***SI appendix*, Figs. S13 and S14, Dataset S1)** signifying pleiotropic effects of certain markers for the VIs. These pleiotropic effects have various associations with developing young tissues, inflorescence, and yield.

### Phenomic prediction can perform similarly to or outperform genomic prediction

Phenomic data (TPP_Multi and TPP_RGB) predicted grain yield as well as genomic data using ridge regression (**Fig. 3**) but different results were observed depending on the cross validation scheme. TPP_RGB contained 35 VIs derived from only RGB bands and Weibull_ CHM belonging to fifteen time points (525 phenomic features) resulting in an accuracy of 0.71; this accuracy was same as the accuracy of 0.71 belonging to GP containing the 153,252 segregating whole genome markers. However, when TPP_Multi, which contains the 89 VIs derived from the multispectral bands and Weibull_CHM belonging to twelve time points (1068 phenomic features), were used in the prediction the yield, prediction accuracy reached up to 0.80; substantially higher than both GP and TPP_RGB supplied for the untested genotype in tested environments schemes (CV2) (**Fig. 3**). Moreover, in the most challenging cross-validation scheme, untested genotypes in untested environment (CV4), GP, TPP_RGB and TPP_Multi performed approximately equally as their prediction accuracies were around 0.50 ± 0.05 (**Fig. 3**). These empirical findings suggest, for the first time, that increasing temporal as well as spectral information can be used to predict fitness substantially better than genomic prediction. This also suggests that temporal and continuous phenomic data can be better predictors than discrete genomic data in prediction and selection of high yielding genotypes. In the only two previous phenomic prediction studies reported to date, (2) used 3,076 NIRS bands at a single timepoint, while (40) used 1,050 NIRS bands on grain samples. (40) then showed these NIRS bands outperformed genomic selection which used 84,259 SNP markers in wheat. Overall, phenomic selection is an emerging approach that may remove the cost of genotyping each year that is required by genomic prediction/selection. Adding a temporal component into phenomic prediction has innumerable known and yet to be discovered advantages.

In summary, this study demonstrated the predictive capability of phenomic data for complex traits in maize, yielding as much as genomic markers frequently applied in plant selection over the past 20 years. UAS surveys over the experimental field plots supplied temporal traits as predictors to facilitate the selection of untested genotypes in untested environments. Growing more plants and measuring them accurately are critical steps to drive effectiveness of selection intensity and accuracy resulting in higher genetic gain over time. This study exemplified that screening more plants and measuring them thanks to repetitive UAV flights across plant growth may results in greater genetic gain than genomic selection when phenomic prediction/selection is applied routinely.

## Acknowledgements

The authors acknowledge Dr. Sorin Popescu and Dr. Lonesome Malambo for the implementing RGB flights, and Dr. Dale Cope for implementing multispectral flights over the maize breeding nurseries multiple times and across two years. The authors thank the Genome to Field project, which collaborated with many researchers at many research institutions and made this research possible. The authors also would like to acknowledge David Rooney, Jacob Pekar, Stephan Labar for their technical field support, and graduate and undergraduate/high school students for their dedicated work in the field. A.A was supported by a fellowship from Republic of Turkey, Ministry of National Education and Ministry of Agriculture and Forestry.

## Author contributions

A.A.: conceptualization, data curation, formal analysis, investigation, methodology, original draft, review and editing (lead), supervision, validation, visualization; S.C.M.: conceptualization, funding acquisition, methodology, project administration, supervision, resources, review and editing (supporting); S.L.A.: conceptualization, data curation, formal analysis, investigation, methodology, review and editing (supporting)

## Supporting Information

Adak *et al*. Temporal unoccupied aerial system phenomic predictions can outperform genomic predictions

## SI Materials and Methods

### Phenomic data extraction pipeline

Detailed function settings of R/UAStools::plotshpcreate were set as follows: (i) *nrowplot* was set 2 since two consecutive row plots represent the one hybrid pedigree, *multirowind* was also set TRUE (T) to define two consecutive row plots indicates one pedigree; (ii) dimension of each polygon was defined by setting the functions of *rangespc* and *rowspc* as 7.62 and 0.76 meters respectively; (iii) buffer polygon was obtained by removing the alley distances from left, right, top and bottom sides using *rangebuf* and *rowbuf* functions; buffer polygon was obtained by setting *rangebuf* (for top and bottom sides) and *rowbuf* (for left and right sides) as 0.61 and 0.05 meters respectively. Buffer polygons covering each plot were used as shape files in data extraction pipelines to obtain better accuracy since walking alleys surrounding the plots were excluded (https://github.com/andersst91/UAStools/wiki/plotshpcreate.R). As a result, a shape file containing 594 buffer polygons (each contain two row plots) were created with the unique plot number in each. After constructing the shape file, each buffer polygon was visualized with tiff files for each time point in QGIS software (https://qgis.org/en/site) and checked manually; occasionally a small percentage of polygons was required to move slightly around the row plots to make each cover the row plots accurately because of the minor overlap issue for certain region of the mosaicked tiff files.

To extract the VIs, first, the tiff files were clipped into a trial level in QGIS, then the extraction pipeline was applied to each clipped tiff file in R. Extraction pipeline were explained briefly as follows: (i) *aggregate* function was first implemented to each tiff file consistently to reduce the computational time requirement by setting *fact* as 4 [*aggregate(“input tiff file”, fact = 4*]; (ii) soil was removed from the tiff files by using the Hue index in *R/FIELDImageR::fieldMask* function; (1); (iii) additional VIs for both HTP platforms were defined in *R/FIELDImageR::fieldndex* function using the output tiff file of second step; (iv) previously constructed shape files were combined with the output of the third step to obtain the values of each VI for each row plot. The VIs calculated by using RGB bands were extracted from the images in first HTP platform while VIs calculated by using RGB, red edge and NIR bands were extracted from the images in second HTP platform.

To construct the canopy height model (CHM), each 3D point cloud file (.las) was first clipped into trial level then the following steps of the custom batch code was applied to each point cloud to extract the plot based temporal plant height as follows: (i) sorting the clipped point clouds to facilitate further processing steps (LAStools/lasssort.exe); (ii) removing excessively noise points (blunders) located below ground and above canopy (LAStools/lasnoise.exe) of row plots; (iii) using the hierarchical robust interpolation algorithm (HRI) (2) to determine the ground points (FUSION\GroundFilter.exe); (iv) detecting the key points from the ground filter to outline digital terrain model (DTM) (LAStools\lasthin.exe); (v) creating the DTM model using the key points from the previous step (FUSION\GridSurfaceCreate.exe); (vi) generating the canopy surface model by extracting the digital terrain model (output of step v) from the digital surface model (output of step ii) (LAStools\lasheight.exe). Adjusting ‘Z’ values that account for plant height in the canopy surface model, merging with the ESRI shape file to clip the canopy surface model into plots (FUSION/PolyClipData.exe). As a last step, statistical metrics (e.g., plant height values based on different percentiles) for each clipped plot (pedigree row) were calculated (FUSION/CloudMetrics.exe). Predicted CHM for each pedigree by equation 2 (*Eq. 2*) was fit based on the Weibull sigmoidal growth model as follows:

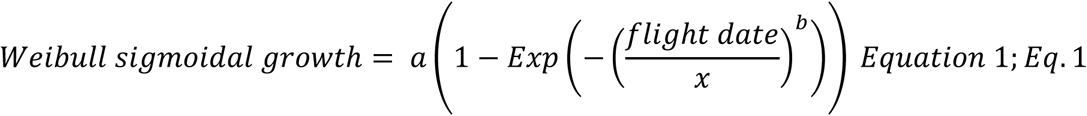

Where, *a* is the asymptote; *flight date* is numeric values as days after planting of each flight date; *x* is the inflection point and *b* is the growth rate. Weibull fit CHM was used in further analysis.

### Experimental design and nested model for phenomic data

Following extraction of plot based temporal vegetation indices and CHM, a nested design predicted the temporal breeding values for each of 280 pedigrees to assess the temporal phenomic data jointly for optimal management (OM) and stressed management (SM, no irrigation, low fertilizer) using the “lmer” function in the “lme4” package in R. Each temporal vegetation index and temporal plant height was modelled for both HTP platform as follows:

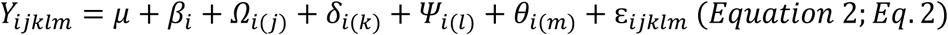

where, *μ* = overall mean; *β*_*i*_ = the random effect of *i*th flight time (as days after planting time, DAP) with 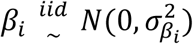, *i* ∈ { 27, 34, 41, 48, 55, 59, 63, 69, 77, 82, 88, 97, 103, 105, 118; rotary-wing with RGB camera HTP platform} and *i* ∈ {27, 34, 52, 60, 70, 73, 88, 105, 112, 118, 132, 144; tuffwing with multispectral camera HTP platform}; *Ω*_*i*(*j*)_ = the random effect of *j*th pedigree (maize hybrid) within the *i*th flight time with 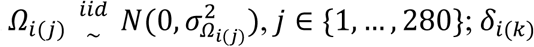, = the random effect of *k*th range within the *i*th flight time with 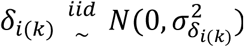, *k* ∈ {1, …, 18}; *Ψ*_*i*(*l*)_= the random effect of *l*th row within the *i*th flight time with 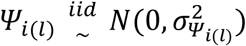, *l* ∈ {1, …, 33}; *θ*_*i*(*m*)_= the random effect of *m*th replication within the *i*th flight time with 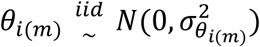, *m* ∈ {1, 2}; ε_*ijklm*_ is pooled error with 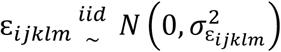.

Temporal repeatability (TR) was calculated using the genotypic variance containing the variation of the trait belonging to all flight times. *Eq. 3* was applied to each vegetation index and canopy height separately.

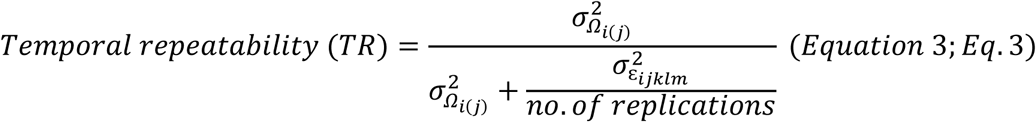

Where, 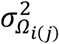 is genotypic variance containing joint genotypic variation occurring across the flights; 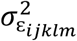 is residual variance containing unexplained error.

Grain yield (GY) was collected from each two adjacent row plots (per hybrid) via a plot combine harvester; days to anthesis (DTA) and silking (DTS) were collected when fifty percent of plots displayed anthesis and silking emergence; manually measured terminal plant height was calculated from the ground to the tip of tassel.

GY, DTA, DTS, PHT were used as predicted variables and modelled according to *Eq. 2* without flight time (denoted as *β* in *Eq. 2*) component as follows:

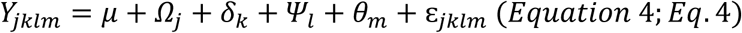

Traditional repeatability was calculated for all cumulative traits (GY, DTA, DTS, PHT) based on

*Eq. 3* with the nested effect by flight time removed (denoted as *β* in *Eq. 2*) as follows:

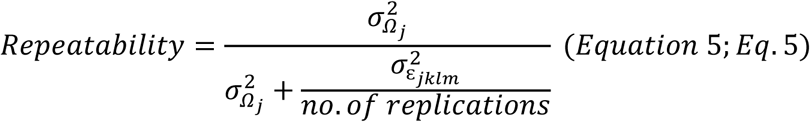

As a result of *Ω*_*i*(*j*)_ component in *Eq*. 2, 35 VIs and Weibull_CHM belonging to fifteen time points in TPP_RGB, and 89 VIs and Weibull_CHM belonging to twelve time points in TPP_Multi were predicted, resulting in 540 and 1080 phenomic data belonging to 280 maize hybrids respectively. Pearson correlation coefficients between each phenotype data at each time point of each temporal trait with GY were calculated using the “*corrplot*” package in R.

### Machine learning based phenomic prediction models

*Caret* package was used in R to run the prediction models. “*Caret::trainControl()*” function was used to set repeated cross validation (*method=“repeatedcv”*) with 10 folds and 3 repeats; this cross validation was used for every model consistently inside the loop. Brief steps of the single loop were explained as follows: (i) partitioning the whole data set as 70 percent training and remainders test data set in TPP_RGB and TPP_Multi phenomic data belonging to optimal (OM) and stress (SM) managements, which were different in each loop, using the “*caret::createDataPartition()*” function, (ii) training the all prediction models using the train data set of OM (tested environment) in the “*caret::train()*” function, (iii) predicting the train data set in OM (cross validation 1; tested genotypes in tested environment; CV1), test data set in OM (cross validation 2; untested genotypes in tested environment; CV2), train data set in SM (cross validation 3; tested genotypes in untested environment; CV3) and test data set in OM (cross validation 4; untested genotypes in untested environment; CV4) using the trained model to obtain the predicted data using the “*caret::predict()*” function for each model, (iv) computing the correlation between actual data and predicted data to evaluate the prediction accuracy 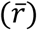 for four cross validation schemes in each model and (v) saving the correlation results along with the R-squared (R^2^), root mean square error (RMSE) and mean absolute error (MAE) as well as the variable importance scores of the predictors belonging to each model in each loop. Number of loops was set to 500. To define the prediction model inside the “*caret::train()*” function, method was set as “*lm*” for the linear model, method was set as “*glmnet*” for elastic net, lasso and ridge models and model was set as “*rf*” for random forest (RF) model separately. To tune the parameters of the elastic net, lasso and ridge regressions, “*alpha*” value was set as 0 for ridge and 1 for lasso regression while sequential numbers between 0 and 1 by ten equal increment numbers were searched to find the best alpha for elastic net regression. Sequential “*lambda*” value between 0 and 1 by ten equal increment numbers were also empirically searched to find the best “*lambda*” values for lasso, ridge, and elastic net regressions. To tune the parameters of RF model, “*ntree*” (number of trees to grow in the model) was set as 1000 while sequential “*mtry*” (number of variables randomly tested as candidates at each split) value between 1 and 50 by five equal increment number were empirically searched to find the best “*mtry*” based on highest accuracy metric of RF model.

### Association mapping for phenomic data

Cumulative AUC were calculated by using the below formula for each pedigree and each VI and Weibull_CHM:

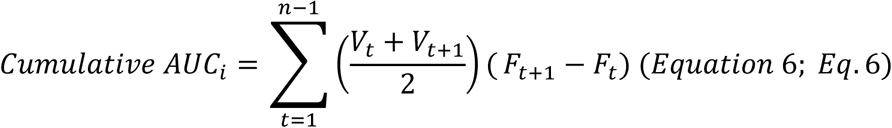

Where, *n* is the number of total observations referring to the fifteen flights times in TPP_RGB, *Cumulative AUC*_*i*_ is the cumulative AUC value based on the total number of flights belonging to *it*h pedigree for each trait, *i* ∈ {1, …, 280}; *V*_*t*_ is the value at *t*th flight time as DAP, *t* ∈ {27, 34, 41, 48, 55, 59, 63, 69, 77, 82, 88, 97, 103, 105, 118}; *F*_*t*_ is the *t*th flight time as number of DAP at which value of interest was taken in first HTP platform.

The imputed ZeaGBSv2.7 with AGPv4 coordinates was used in this study, available in Panzea (https://www.panzea.org/genotypes) and Cyverse (3) platforms. In the genome wide association mapping study, 158 maize hybrids with genotyping by sequencing (GBS) data of their parental lines were available in MaizeGBSv2.7 (4), generated via the method based on digestion the DNA with the *ApeKI* restriction enzyme (5). ZeaGBSv2.7 was called in Tassel 5 software (6). Before association mapping, GBS data of the hybrid maize was created based on following step: (i) heterozygote calls belonging to any parental lines of the hybrids were set as missing, (ii) “*create_hybrid_genotype*” function in Tassel 5 software was used to create the GBS data of hybrid maize by merging the GBS data of parental lines of each hybrid and (iii) polymorphic markers were obtained by filtering the missing data that is more than ten percent and minor allele frequency that is lower than five percent. LD *k*-nearest neighbor algorithm (LD *KNNi* imputation) was implemented to GBS data to impute the missing calls in Tassel software (version 5) (7). Finally, 101,100 polymorphic SNPs (single nucleotide polymorphism) remained and were used in the association mapping analysis.

To control the population structure of the hybrid population, the first five principal components, which explained 49% of total variation, and kinship matrix were used in each model. Bonferroni corrections [− *log*_10_(*p values*) > 6.3; 0.01/(no. of markers)] were considered as threshold in determining the GWAS hits in Manhattan plots, in addition to Bonferroni threshold, false-positive discovery rate was set [− *log*_10_(*p values*) > 5] to detect same loci (if any) that were associated with multiple traits with between the values of [− *log*_10_(*p values*) > 5] and [− *log*_10_(*p values*) > 6.3]. MaizeGBD (http://www.maizegdb.org/) and the Gramene database (http://www.gramene.org) were used to determine corresponding candidate genes of the discovered SNPs and functions of genes. LD decay pattern was investigated in Tassel 5 (LD windows size = 10 markers) and visualized in R for each chromosome separately (**Fig. S1**). Linkage disequilibrium (LD) was visualized using the *LDheatmap* package in R (8) to identify the candidate genes within the LD blocks (*R*^2^ ≥ 0.8) of colocalized SNPs.

### Genomic prediction for phenomic data

153,252 SNPs belonging to 158 maize hybrids were obtained merging the GBS data of their parental lines in Tassel software as described in the “*Association mapping for phenomic data”* section. After obtaining the hybrid GBS, SNPs were filtered if minor allele frequency was lower than 0.01 and missing values were higher than 10 percent per marker resulting in 153,252 SNPs. Missing values of 153,252 SNPs were imputed using the *rrBLUP::Amat()* function in R. Temporal genomic prediction for phenomic data in TPP_RGB was modeled using the rrBLUP package (9) in R as follows:

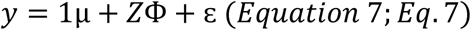

Where, *y* = is the vector (*n* × 1) of single phenotype data of *n* maize hybrids (*n* = is training data set of each loop) belonging to each single time point of each phenotype data in TPP_RGB, 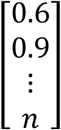 1 = vector of ones that are equal to numbers of *n*,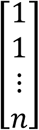; μ = overall mean of training data set; *Z* = the incidence matrix (*n* × *p*) of allelic states of *p* number of SNPs (153,252 SNPs) belonging to *n* number of maize hybrids,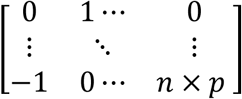; Ф = vector of calculated SNP effects (*p* × 1), 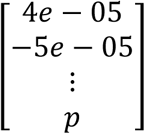; ε = vector of random residuals. RR-BLUP assumes 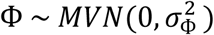 indicating that marker effects are normally distributed with equal marker variance 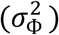 throughout the genome. Genomic prediction was modelled using training data accounting for seventy percent of total data while the remaining thirty percent data was used as validation data. The genomic prediction model (*Equation* 7, *Eq*. 7) evaluated 500 iterations applied to each phenotype of 158 maize hybrids belonging to each VI and Weibull_CHM at fifteen time points (totally 540 phenotype data) in TPP_RGB; base R function called “*sample()*”was used to randomly determine training and test data set in each iteration. During the prediction, the same training and test data set for each phenotype of each trait at each time point was needed to obtain fair comparison of the genomic prediction accuracy. Genomic prediction accuracy was calculated based on correlation results between the genetic estimated breeding values and the true breeding value of test data set in each iteration.

### Phenomic prediction versus genomic prediction

118 maize hybrids whose parental lines had GBS info in MaizeGBSv2.7 (4) and grown in optimal (OM) and stressed (SM) managements were used. In genomic prediction, GBS data (GP) containing 153.252 SNPs that was described in the “*Genomic prediction for phenomic data”* was used to predict grain yield (GY) using rrBLUP package in R (9). TPP_RGB containing 540 phenomic data, and TPP_Multi containing 1080 phenomic data were used to predict GY using the ridge regression in the *caret* package in R. Prediction accuracy was obtained from 1000 bootstraps for each model where the same training and test data set were used for genomic and phenomic prediction within each bootstrap. Genomic prediction and phenomic prediction steps were explained in the *“Genomic prediction for phenomic data”* and *“Machine learning based phenomic prediction models”* respectively. Four cross validation schemes, which were explained in *“Machine learning based phenomic prediction models”*, were applied in this section as well to compare the prediction accuracies of two phenomic predictions and genomic prediction.

**Table S1.**
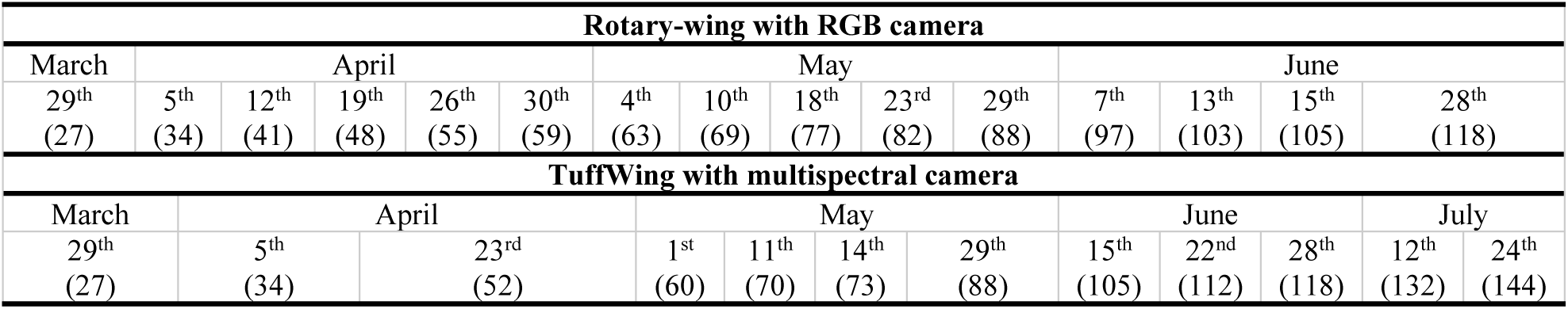
shows the calendar days between March to July, 2017 containing the flight dates for rotary-wing UAS with RGB camera (above) and TuffWing UAS with multispectral camera (below) high throughput phenotyping platforms. Flight dates were shown under the months with corresponding days after planting times in parenthesis.

**Table S2.** shows the vegetation indices used and their formulas along with references.

| Vegetation index | Ratios | References |
| --- | --- | --- |
| VIs derived from RGB bands |  |  |
| Blue chromatic coordinate index (BCC) | $\frac{B}{R + G + B}$ | (10) |
| Blue green pigment index (BGI) | $\frac{B}{G}$ | (11) |
| Brightness index (BI) | $\text{sqrt}(\frac{R^2 + G^2 + B^2}{3})$ | (12) |
| Color index of vegetation extraction (CIVE) | $0.441R - 0.811G + 0.385B + 18.78745$ | (13) |
| Combined indices 1 (COM1) | $EXG + CIVE + EXGR + VEG$ | (14) |
| Combined indices 2 (COM2) | $0.36EXG + 0.47CIVE + 0.17VEG$ | (15) |
| Additional blue index (EBI) | $\frac{B - G}{B - R}$ | (16) |
| Additional green index (EGI) | $\frac{G - R}{R - B}$ | (16) |
| Green-red index (ERI) | $\frac{R - G}{R - B}$ | (16) |
| Excessive green (EXG) | $2G - R - B$ | (10) |
| Normalized Excess green index (EXG2) | $\frac{2G - R - B}{G + R + B}$ | (10) |
| Excess green minus excess red index (EXGR) | $3G - 2.4R - B$ | (17) |
| Excessive red (EXR) | $1.4R - G$ | (18) |
| Green minus blue index (G-B) | $G - B$ | (10) |
| Green minus red index (G-R) | $G - R$ | (10) |
| Green blue simple ratio index (G/B) | $\frac{G}{B}$ | (10) |
| Green red simple ratio index (G/R) | $\frac{G}{R}$ | (10) |
| Green chromatic coordinate index (GCC) | $\frac{G}{R + G + B}$ | (10) |
| Green leaf index (GLI) | $\frac{2G - R - B}{2G + R + B}$ | (19) |
| Modified excess green index (MEXG) | $1.262G - 0.884R - 0.311B$ | (20) |
| Modified green red index (MGVRI) | $\frac{G^2 - R^2}{G^2 + R^2}$ | (21) |
| Normalized difference index (NDI) | $128 * \left[ \left( \frac{(G - R)}{(G + R)} \right) + 1 \right]$ | (17) |
| Normalized difference red blue index (NDRBI) | $\frac{R - B}{R + B}$ | (16) |
| Normalized green-blue difference index (NGBDI) | $\frac{G - B}{G + B}$ | (22) |
| Normalized green red difference index (NGRDI) | $\frac{G - R}{G + R}$ | (23) |
| Red minus blue index (R-B) | $R - B$ | (10) |
| Red blue simple ratio index (R/B) | $\frac{R}{B}$ | (10) |
| Red chromatic coordinate index (RCC) | $\frac{R}{R + G + B}$ | (10) |
| Red green blue index (RGBVI) | $\frac{G^2 - R * B}{G^2 + R * B}$ | (21) |
| Triangular greenery index (TGI) | $G - (0.39R - 0.69B)$ | (24) |
| Visible atmospherically resistant index (VARI) | $\frac{G - R}{G + R - B}$ | (25) |
| Vegetative (VEG) | $\frac{G}{R^{0.667} * B^{0.334}}$ | (26) |
| **Vis derived from multispectral bands (RGB, red-edge and NIR bands)** |  |  |
| Modified chlorophyll absorption in reflectance index 1(MCARI1) | $[(NIR - RE) - 0.2 * (NIR - G)] * \left( \frac{NIR}{RE} \right)$ | (27) |
| Modified chlorophyll absorption in reflectance index 2(MCARI2) | $\frac{1.5(NIR - RE) - 1.3(NIR - G)}{\sqrt{(2NIR + 1)^2 - (6NIR - 5\sqrt{RE})} - 0.5}$ | (28) |
| Chlorophyll vegetation index-green (CIG) | $\frac{NIR}{G} - 1$ | (29) |
| Chlorophyll vegetation index-red edge (CIRE) | $\frac{NIR}{RE} - 1$ | (29) |
| Chlorophyll vegetation index (CVI) | $\frac{NIR * R}{G^2}$ | (30) |
| Difference vegetation index (DVI) | $NIR - RE$ | (23) |
| Enhanced normalized difference vegetation index (ENDVI) | $\frac{NIR + G - 2B}{NIR + G + 2B}$ | Maxmax 2015<br>(http://www.maxmax.com/endvi.htm) |
| Enhanced vegetation index (EVI) | $\frac{2.5(NIR - R)}{NIR + 6R - 7.5B + 1}$ | (31) |
| Green difference vegetation index (GDVI) | $NIR - G$ | (23) |
| Green infrared percentage vegetation index (GIPVI) | $\frac{NIR}{NIR + G}$ | (32) |
| Green normalized difference vegetation index (GNDVI) | $\frac{NIR - G}{NIR + G}$ | (33) |
| Green optimal soil adjusted vegetation index (GOSAVI) | $\frac{(1 + 0.16)(NIR - G)}{NIR + G + 0.16}$ | (34) |
| Green re-normalized different vegetation index (GRDVI) | $\frac{NIR - G}{\sqrt{NIR + G}}$ | (35) |
| Green ratio vegetation index (GRVI) | $\frac{NIR}{G}$ | (36) |
| Green soil adjusted vegetation index (GSAVI) | $1.5 \left( \frac{NIR - G}{NIR + G + 0.5} \right)$ | (37) |
| Green wide dynamic range vegetation index (GWDRVI) | $\frac{0.12NIR - G}{0.12NIR + G}$ | (38) |
| Modified double difference index (MDD) | $(NIR - RE) - (RE - G)$ | (39) |
| Modified GSAVI (MGSAVI) | $0.5[2NIR + 1 - \sqrt{(2NIR + 1)^2 - 8(NIR - G)}]$ | (40) |
| Modified normalized difference index (MNDI) | $(NIR - RE)/(NIR - G)$ | (41) |
| Modified normalized difference red edge (MNDRE) | $\frac{[NIR - (RE - 2G)]}{[NIR + (RE - 2G)]}$ | (42) |
| Modified RESAVI (MRESAVI) | $0.5 [2NIR + 1 - \sqrt{(2NIR + 1)^2 - 8(NIR - RE)}]$ | (40) |
| Modified RETVI (MRETVI) | $1.2[1.2(NIR - G) - 2.5(RE - G)]$ | (28) |
| Modified simple ratio (MSR) | $\frac{\left(\frac{NIR}{R} - 1\right)}{\sqrt{\left(\frac{NIR}{R} + 1\right)}}$ | (43) |
| Modified green simple ratio (MSR_G) | $\frac{\left(\frac{NIR}{G} - 1\right)}{\sqrt{\left(\frac{NIR}{G} + 1\right)}}$ | (43) |
| Modified green simple ratio (MSR_RE) | $\frac{\left(\frac{NIR}{RE} - 1\right)}{\sqrt{\left(\frac{NIR}{RE} + 1\right)}}$ | (43) |
| Modified transformed CARI (MTCARI) | $3 \left[ (NIR - RE) - 0.2 \left( NIR - G \right) \left( \frac{NIR}{RE} \right) \right]$ | (44) |
| Normalized difference red edge (NDRE) | $\frac{NIR - RE}{NIR + RE}$ | (45) |
| Normalized difference vegetation index (NDVI) | $\frac{NIR - R}{NIR + R}$ | (23) |
| Normalized NIR index (NNIR) | $\frac{NIR}{NIR + RE + G}$ | (37) |
| Normalized red edge index (NREI) | $\frac{RE}{NIR + RE + G}$ | (37) |
| Normalized green index (NGI) | $\frac{G}{NIR + RE + G}$ | (37) |
| Optimized soil-adjusted vegetation index (OSAVI) | $\frac{NIR - R}{NIR + R + 0.16}$ | (34) |
| Plant senescence reflectance index (PSRI) | $\frac{R - G}{RE}$ | (46) |
| Red edge green difference vegetation index (REGDVI) | $RE - G$ | (23) |
| Red edge GNDVI (REGNDVI) | $\frac{RE - G}{RE + G}$ | (33) |
| Red edge green ratio vegetation index (REGRVI) | $\frac{RE}{G}$ | (47) |
| Red edge optimal soil adjusted vegetation index (REOSAVI) | $\frac{(1 + 0.16)(NIR - RE)}{NIR + RE + 0.16}$ | (34) |
| Renormalized difference vegetation index (RERDVI) | $\frac{NIR - RE}{\sqrt{NIR + RE}}$ | (35) |
| Red edge soil adjusted vegetation index (RESAVI) | $1.5 \left[ \frac{NIR - RE}{NIR + RE + 0.5} \right]$ | (37) |
| Red edge transformed vegetation index (RETVI) | $0.5[120(NIR - G) - 200(RE - G)]$ | (48) |
| Red edge wide dynamic range vegetation index (REWDRVI) | $\frac{0.12NIR - RE}{0.12NIR + RE}$ | (38) |
| Ratio vegetation index (RVI) | $\frac{NIR}{R}$ | (49) |
| Soil-adjusted vegetation index (SAVI) | $\frac{1.5(NIR - R)}{NIR + R + 0.5}$ | (50) |
| Triangular vegetation index (TVI) | $0.5[120(NIR - G) - 200(R - G)]$ | (48) |
| Optimized vegetation index 1 (VI<sub>opt1</sub>) | $100(\ln NIR - \ln RE)$ | (51) |
| Transformed Normalized Vegetation Index (TNDVI) | $\text{sqrt} \left( \frac{NIR - R}{NIR + R} + 0.5 \right)$ | (52) |
| Modified Nonlinear Index (MNLi) | $\frac{1.5(NIR^2 - R)}{NIR^2 + R + 0.5}$ | (53) |
| Red Edge Simple Ratio (RESR) | $\frac{RE}{R}$ | (54) |
| Red edge normalized difference vegetation index (RENDVI) | $\frac{RE - R}{RE + R}$ | (55) |
| Normalized NIR index2 (NNIR2) | $\frac{NIR}{NIR + RE + R}$ | (37) |
| Normalized red edge index2 (NREI2) | $\frac{RE}{NIR + RE + R}$ | (37) |
| Normalized red index (NRI) | $\frac{R}{NIR + RE + R}$ | (37) |

R, G, B, RE and NIR represent the red, green, blue, red-edge and near infrared reflectance bands respectively. Red, green, blue, red edge and NIR reflectance bands were also used in this study singly.

## SI Results

### Genome wide association mapping results

Two close large effect locus 39,906,105 bp (*chr2_2*) and 39,906,547 bp (*chr2_3*) genomic locations was discovered for EXG2, GLI, RGBVI, VEG, Blue, GCC and NDRBI by all three GWAS models consistently and explained between 6 and 51 percent phenotypic variation depending on the traits and GWAS models (**Dataset S1**). *GRMZM2G129493* (chr2: 39906034 to 39907044 bp) candidate gene covers *chr2_2* (39906105 bp) and *chr2_3* (39906547 bp) GWAS peaks in its genomic region; known as polygalacturonase-inhibiting proteins (*PGIPs*) these encode plant defense related proteins (56). Another candidate gene (∼2kb away from *chr2_2 and chr2_3*), *GRMZM2G362362* (chr2: 39893309 to 39904028), belongs to a family of glycoside hydrolases that hydrolase the glycosidic bonds in polysaccharide in cell wall (57).

The 50,705,765 bp (*chr2_4*) genomic location in chromosome 2 was discovered for TGI, Blue, BI, Green and Red by all three models consistently and explained 6 to 7 percent variation depending on the traits and models (**Dataset S1**). GRMZM2G018059 (chr2: 50696420 to 50706825) candidate gene contains the *chr2_4* GWAS peak in its genomic region and its function is related to U-box domain-containing protein kinase family protein that was discovered in previous association mapping studies as drought responsive genes (58, 59) as well as for yield related traits in maize (60).

203,544,095 bp (*chr4_1*) genomic location in chromosome 4 was discovered for TGI, BCC, BGI, G/B, G/R, GCC, NDRBI, NGBDI, R/B, COM1, EXG2, GLI and RGBVI and explained between 3 to 44 percent variation depending on the traits and models (**Dataset S1**). The *GRMZM2G001541* (chr4: 203544095 to 203547230 bp) candidate gene is closest \and ∼30 base pairs away from the *chr4_1* (203544095 bp). The homolog of this gene in *Arabidopsis* is responsible for encoding the inflorescence and root apices receptor-like kinase (IRK) protein that is crucial for maintaining the differentiation of meristem (61). *GRMZM2G001541* has been discovered by meta-QTL and GWAS analysis consistently and found to be highly expressed in developing tissues (e.g. primordia, developing leaves and ear) closely related to inflorescence development (62) influencing yield performance directly in maize (63). *GRMZM2G001541* governs the expression level of *Unbranched3* (UB3), which regulates the quantitative variation of kernel row number in maize (64).

**Figure S1.**
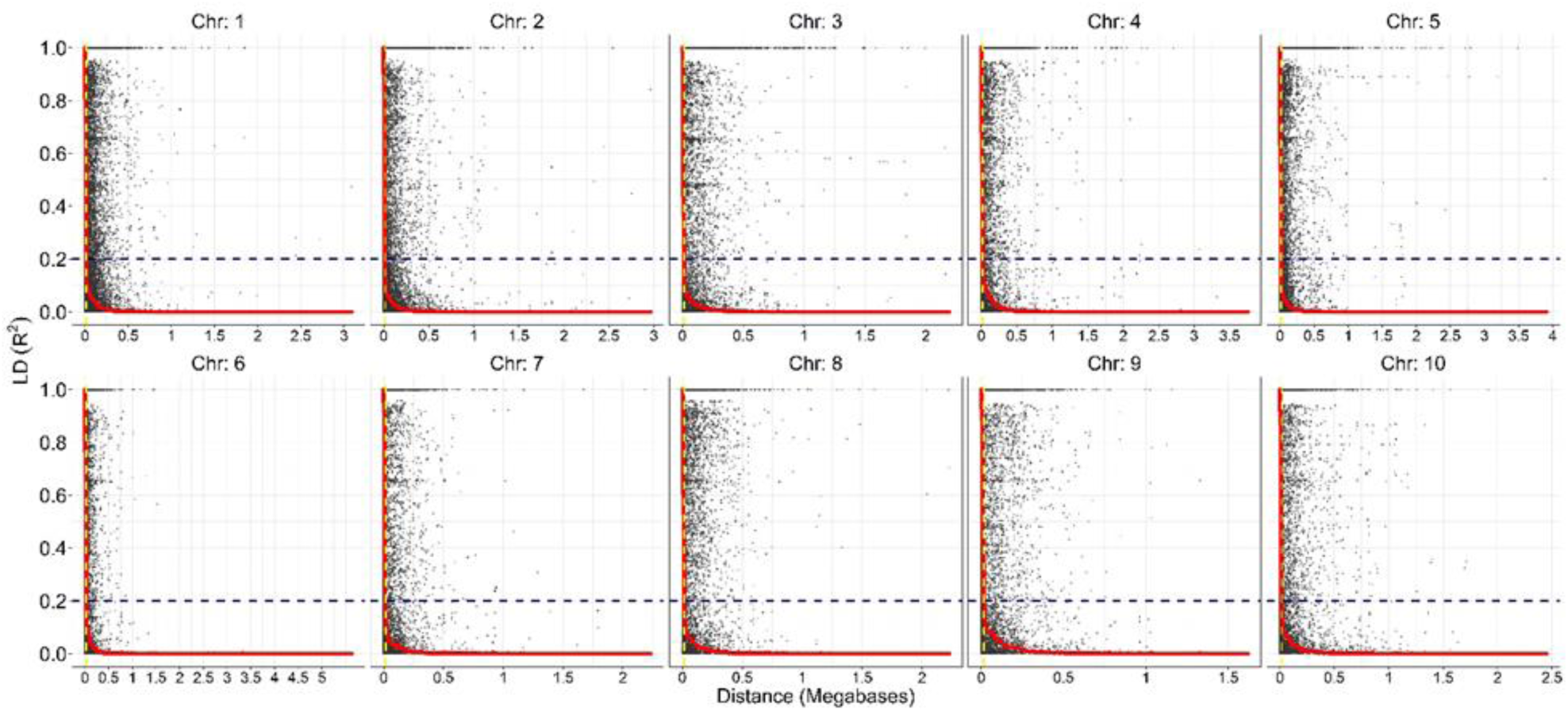
shows linkage disequilibrium decay (LD) patterns for each chromosome. The Y axis represents the R^2^ while X axis shows the distance as Megabases. Horizontal dashed line shows the 0.2 R^2^ LD while the vertical yellow color dashed line shows the 15 kilo base pair (kb) threshold in each chromosome. LD decay was found to be rapid for all chromosome and changing between 1 to 15 kb.

**Figure S2.**
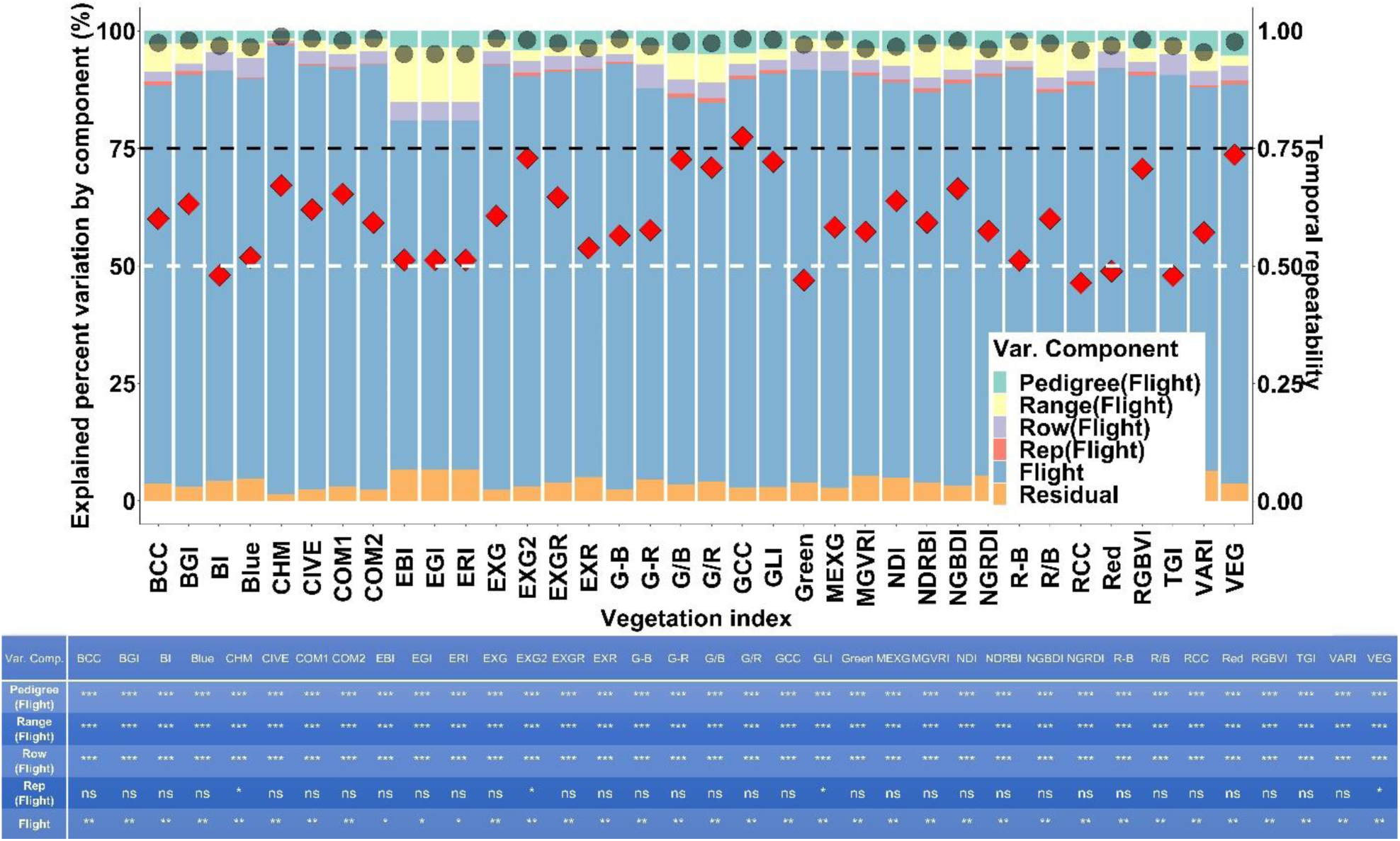
stacked bar plots show the explained percent variation by each component in *Eq. 2* for each temporal trait in TPP_RGB. Left y axis corresponds to the explained percent variation of the components in the stacked bar plots while the right y axis shows the temporal repeatability (red diamonds calculated by Eq 3) and R^2^ values (black round symbols). Gray and black horizontal dashed lines represent the values of 0.50 and 0.75 where most of the temporal repeatability values of temporal traits accumulated. The table below shows the significance values of each component in *Eq. 2* for each temporal trait; ***, **, * are the 0.001, 0.01 and 0.05 significance levels respectively while ns is not statistically significant.

**Figure S3.**
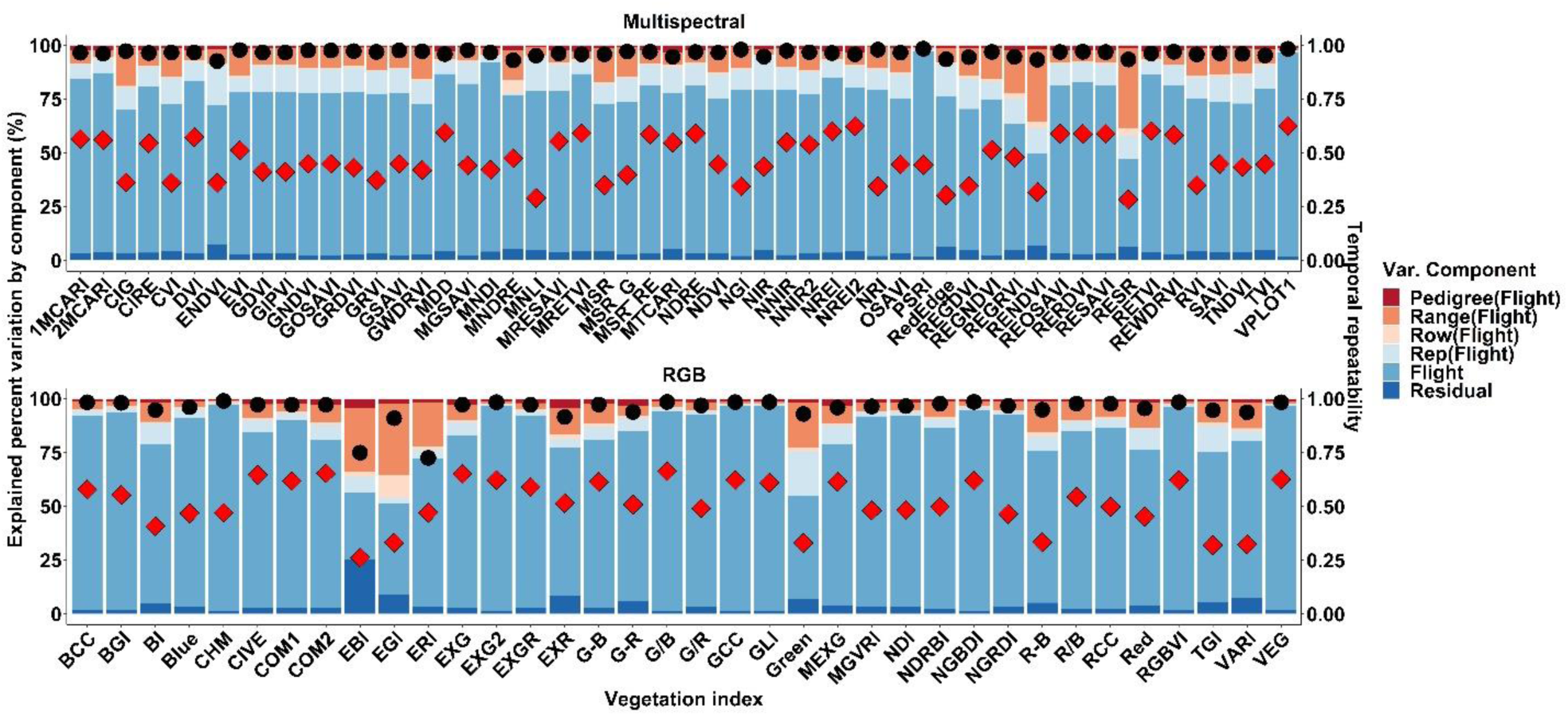
stacked bar plots represent the explained percent variation by each component in *Eq. 2* for each temporal trait in TPP_Multi. Left y axis corresponds to the explained percent variation of the components while right y axis shows the temporal repeatability (red diamonds calculated by *Eq. 3*) and R^2^ values (black round symbols).

**Figure S4.**
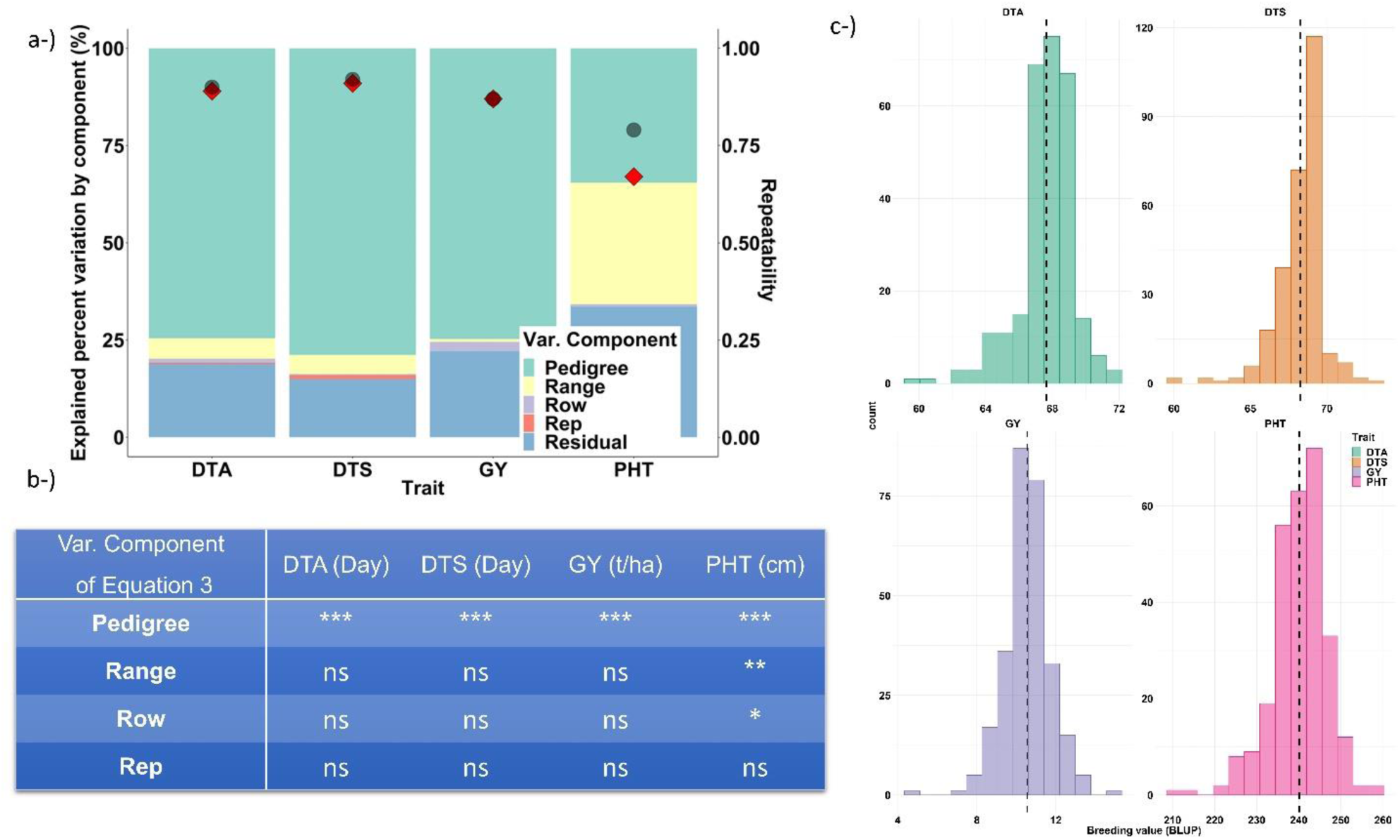
a-) stacked bar plots shows the explained percent variation by each component in *Eq. 4* for each trait. Left y axis represents the explained percent variation of the components while left y axis shows the temporal repeatability (red diamonds calculated by *Eq. 5*) and Rsquared values (black round symbols). b-) The table shows the significance values of each component in *Eq. 4* for each temporal trait; ***, **, * are the 0.001, 0.01 and 0.05 significance levels respectively while ns is not statistically significant. c-) shows the histograms of the breeding values of each trait with their means represented by vertical black lines.

**Figure S5.**
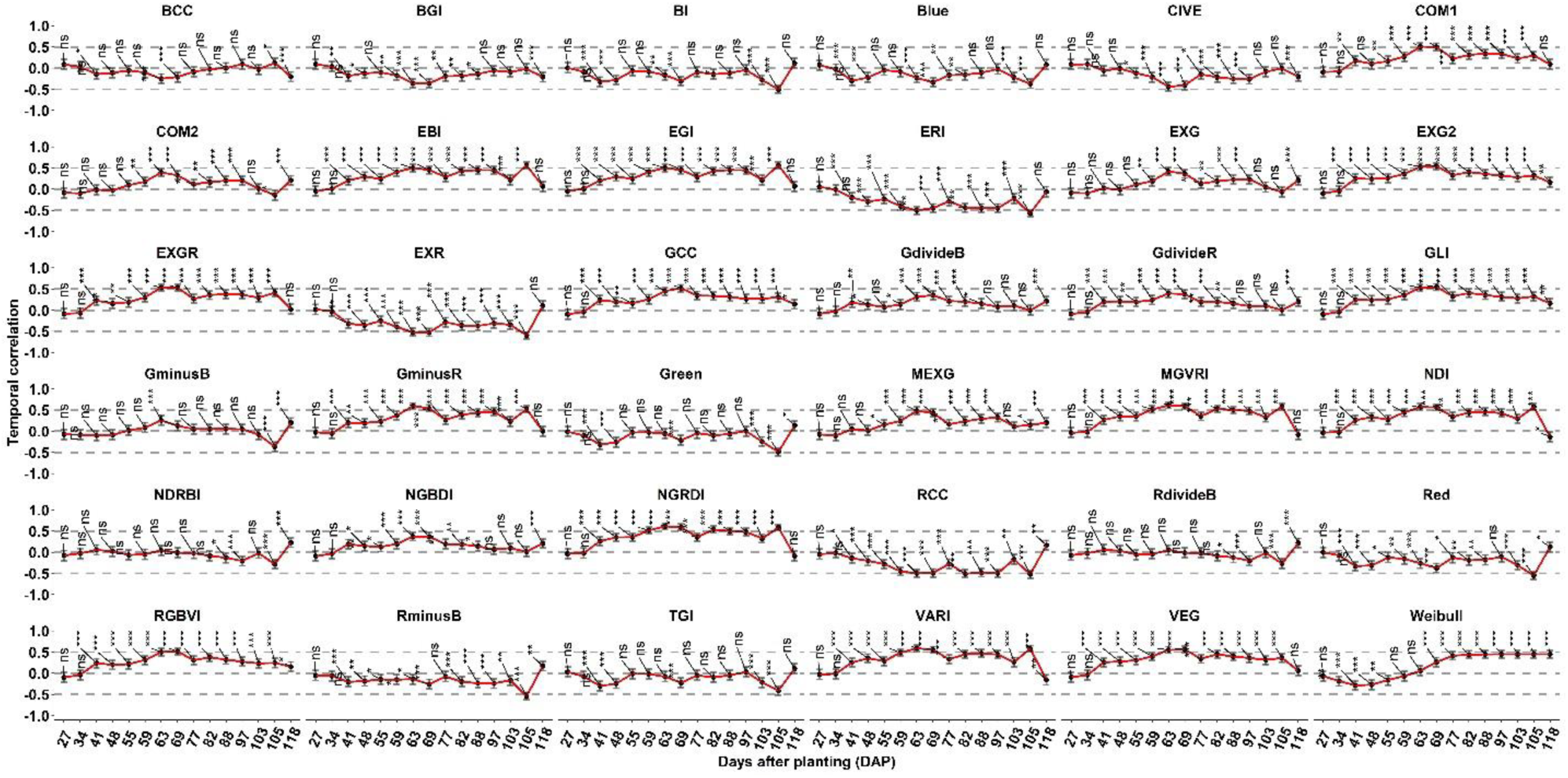
Pearson correlation coefficients between the phenotype values at each flight time point of the VIs in TPP_RGB. Whiskers shows upper and lower confident intervals of temporal correlation based on 95 percent confidence level. Above, middle, and below dashed lines represent the 0.5, 0 and -0.5 temporal correlation values. ***, **, * are the 0.001, 0.01 and 0.05 significance levels respectively while ns is not statistically significant.

**Figure S6.**
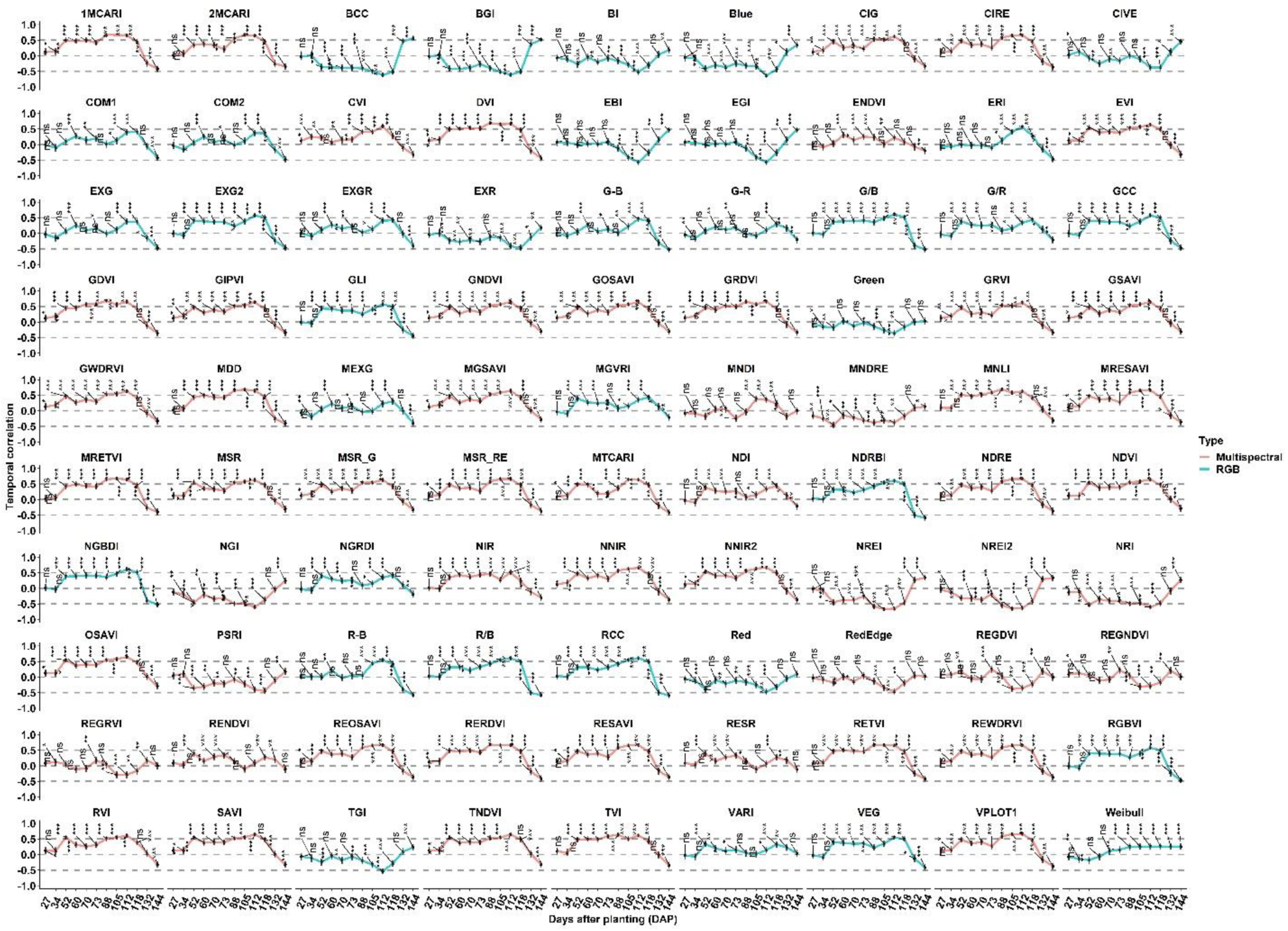
Pearson correlation between the phenotype values at each flight time point of the VIs in TPP_Multi. The title separated vegetation indices according to their derivation from multispectral bands and RGB bands. Whiskers show upper and lower confidence intervals of temporal correlation based on 95 percent confidence level. Above, middle, and below dashed lines represent the 0.5, 0 and -0.5 temporal correlation values. ***, **, * are the 0.001, 0.01 and 0.05 significance levels respectively; ns is not statistically significant.

**Figure S7.**
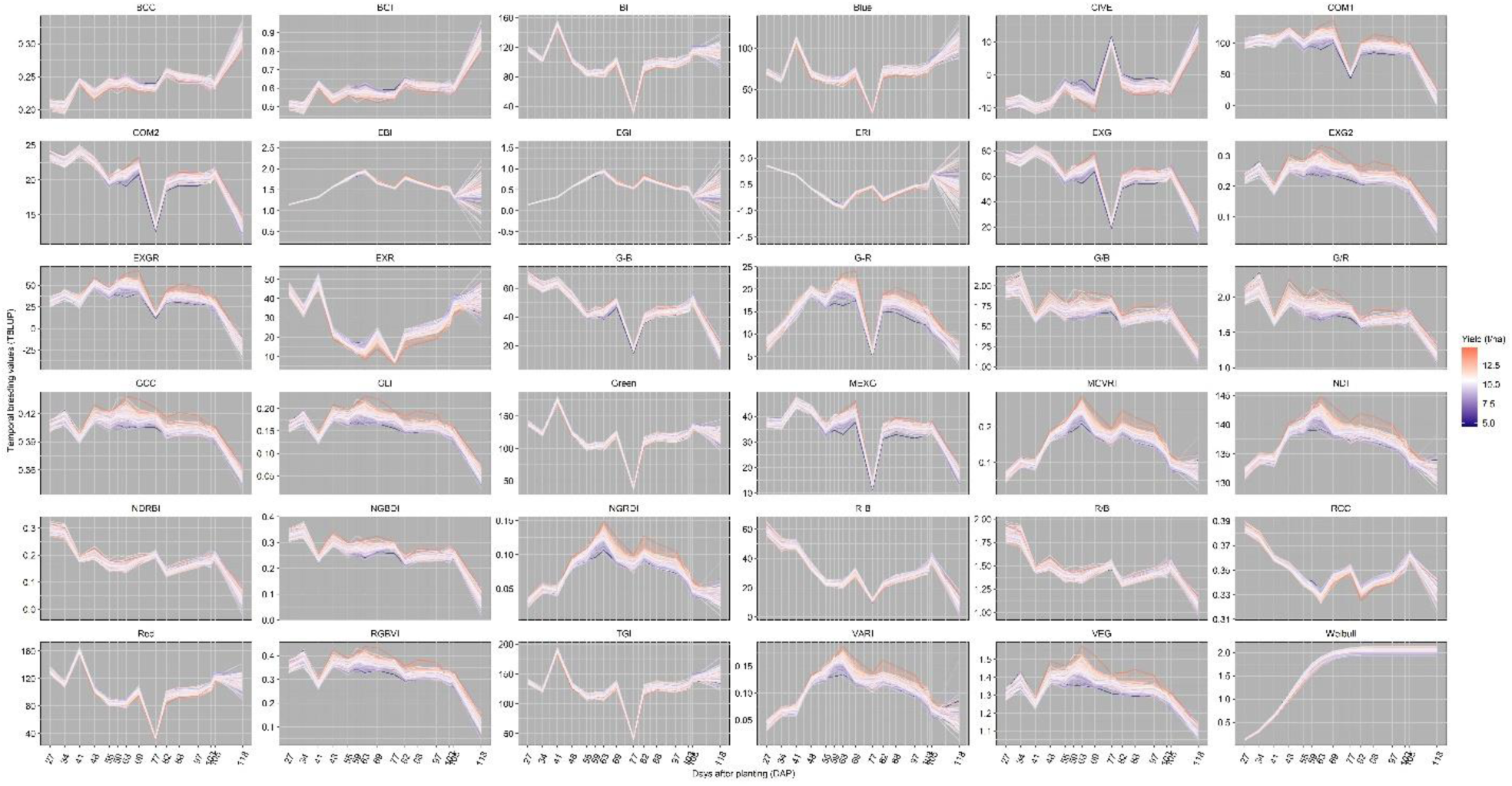
temporal pedigree values of 280 maize hybrids predicted by *Eq. 2* belonging to each vegetation indices and Weibull_CHM in TPP_RGB. Each hybrid was colored according to their yield values that are low, average, and high yield values representing blue, white, and red respectively in the heatmap scale. Average yield was 10.5 t/ha.

**Figure S8.**
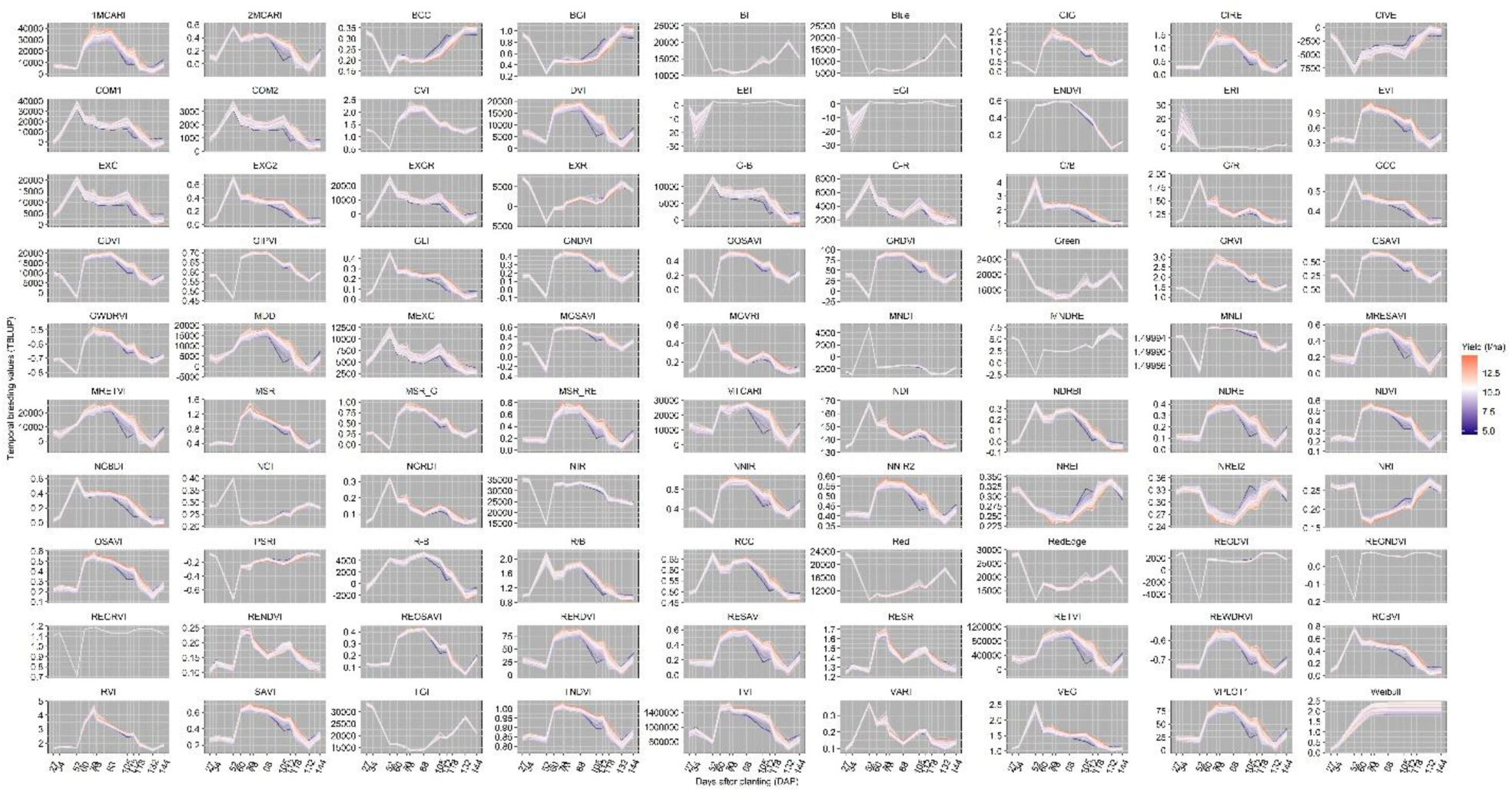
temporal pedigree values of 280 maize hybrids predicted by *Eq. 2* belonging to each vegetation indices in TPP_Multi. Each hybrid was colored according to their yield values that are low, average, and high yield values representing blue, white, and red respectively in the heatmap scale. Average yield was 10.5 t/ha.

**Figure S9.**
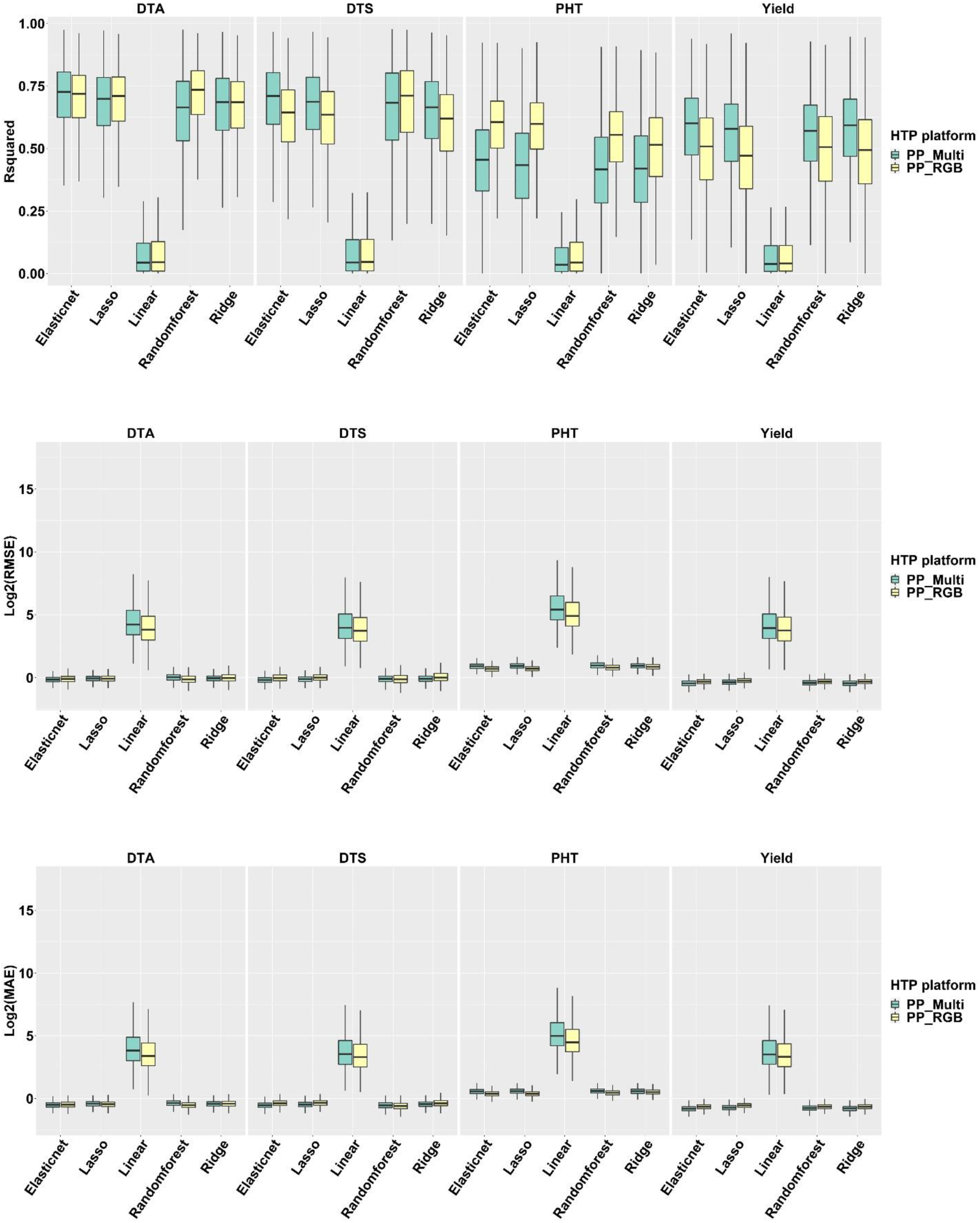
R-squared, root mean square error (RMSE), and mean absolute error (MAE) values from top to bottom belonging to each model (on the x axis) and each predicted variable (from left to right). Log_2_() transformation was applied to RMSE and MAE values to show the excessive values belonging to linear model.

**Figure S10.**
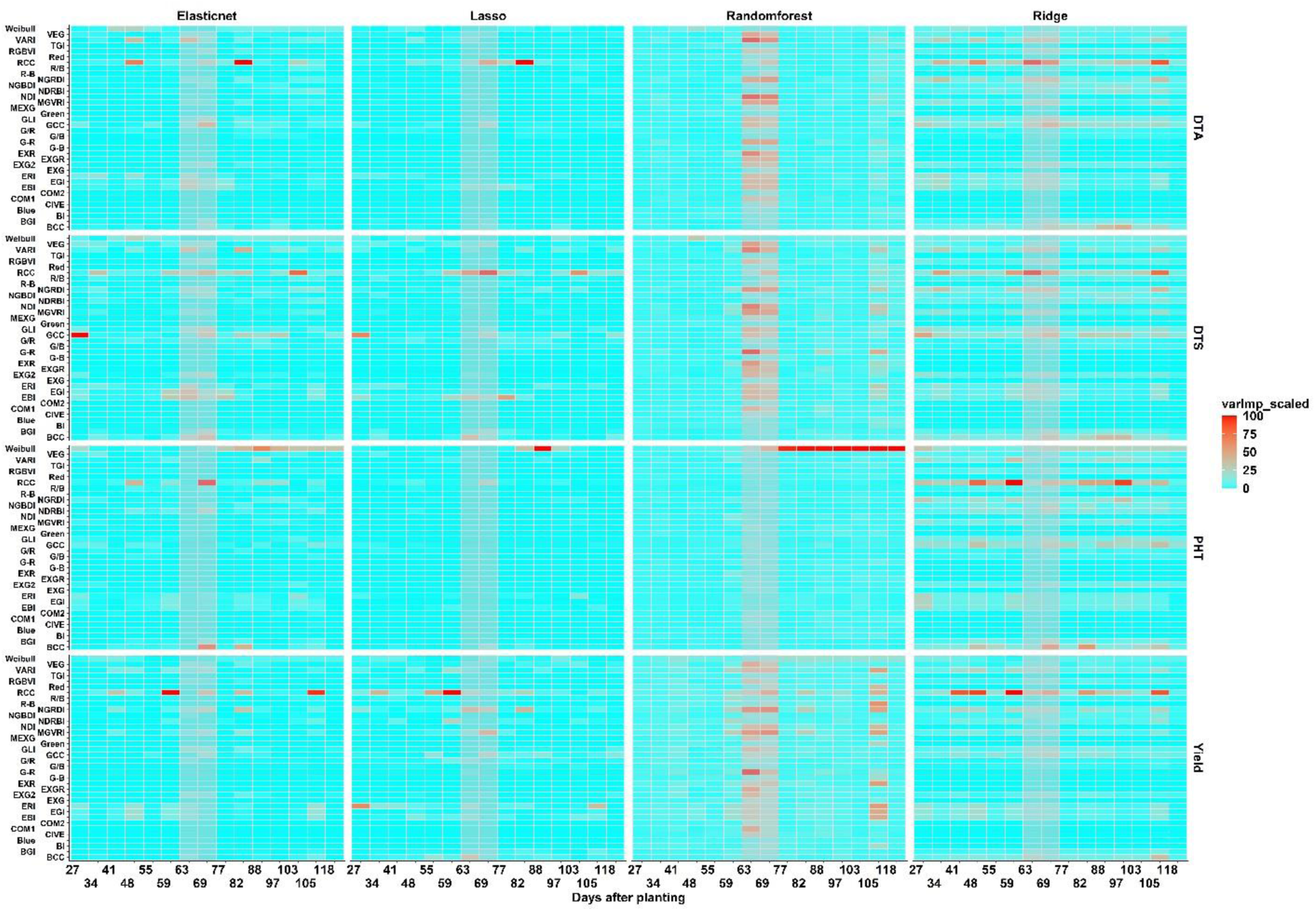
Variable importance scores belonging to each predicted variable (from top to bottom) and model (from left to right) when TPP_RGB phenomic data was used. X axis shows the VIs as well as Weibull_CHM and X axis shows the flight dates as days after planting times. The highlighted grey columns correspond to the range of flowering dates in this population.

**Figure S11.**
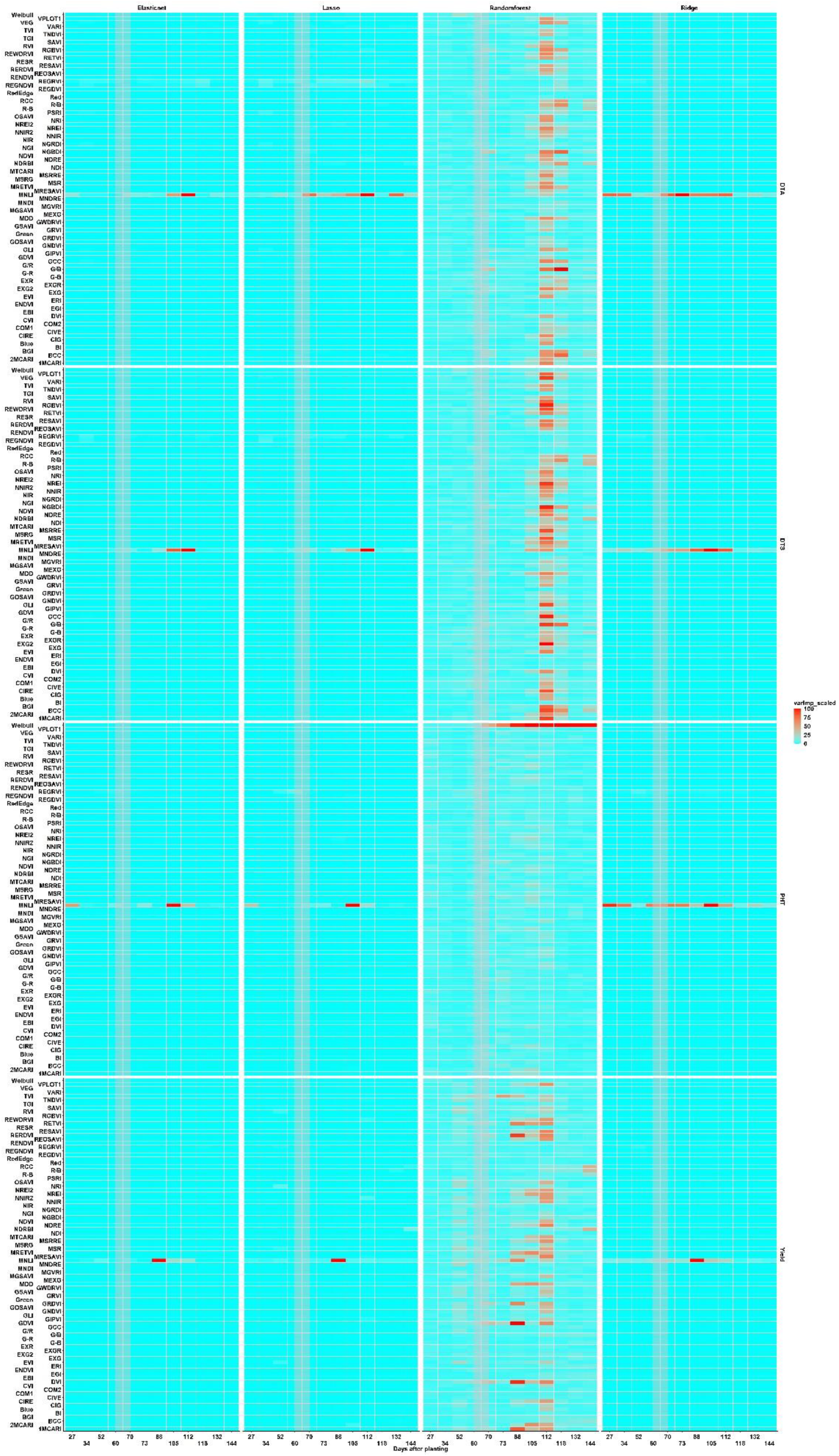
Variable importance scores belonging to each predicted variable (from top to bottom) and model (from left to right) when TPP_Multi phenomic data was used. X axis shows the VIs as well as Weibull _CHM and X axis shows the flight dates as days after planting times. The highlighted grey columns correspond to the range of flowering dates in this population.

**Figure S12.**
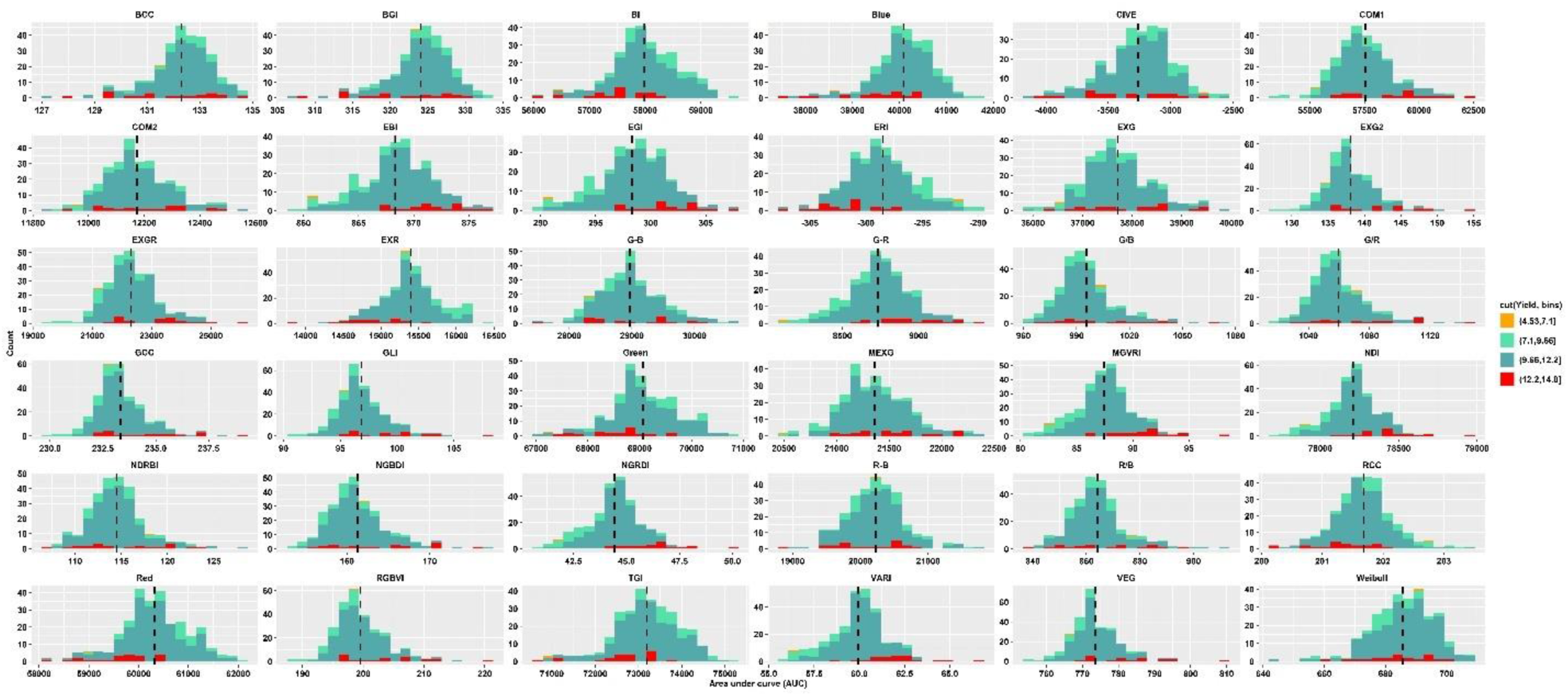
Histograms of area under curve value of each VI and Weibull _CHM for each genotype calculated by *Eq. 6* in TPP_RGB phenomic data. Area under curve values were divided into four equal bin categories based on yield value as shown in the legend and each histogram was colored based on these bins. Vertical dashed lines in each histogram show the mean values.

**Figure S13.**
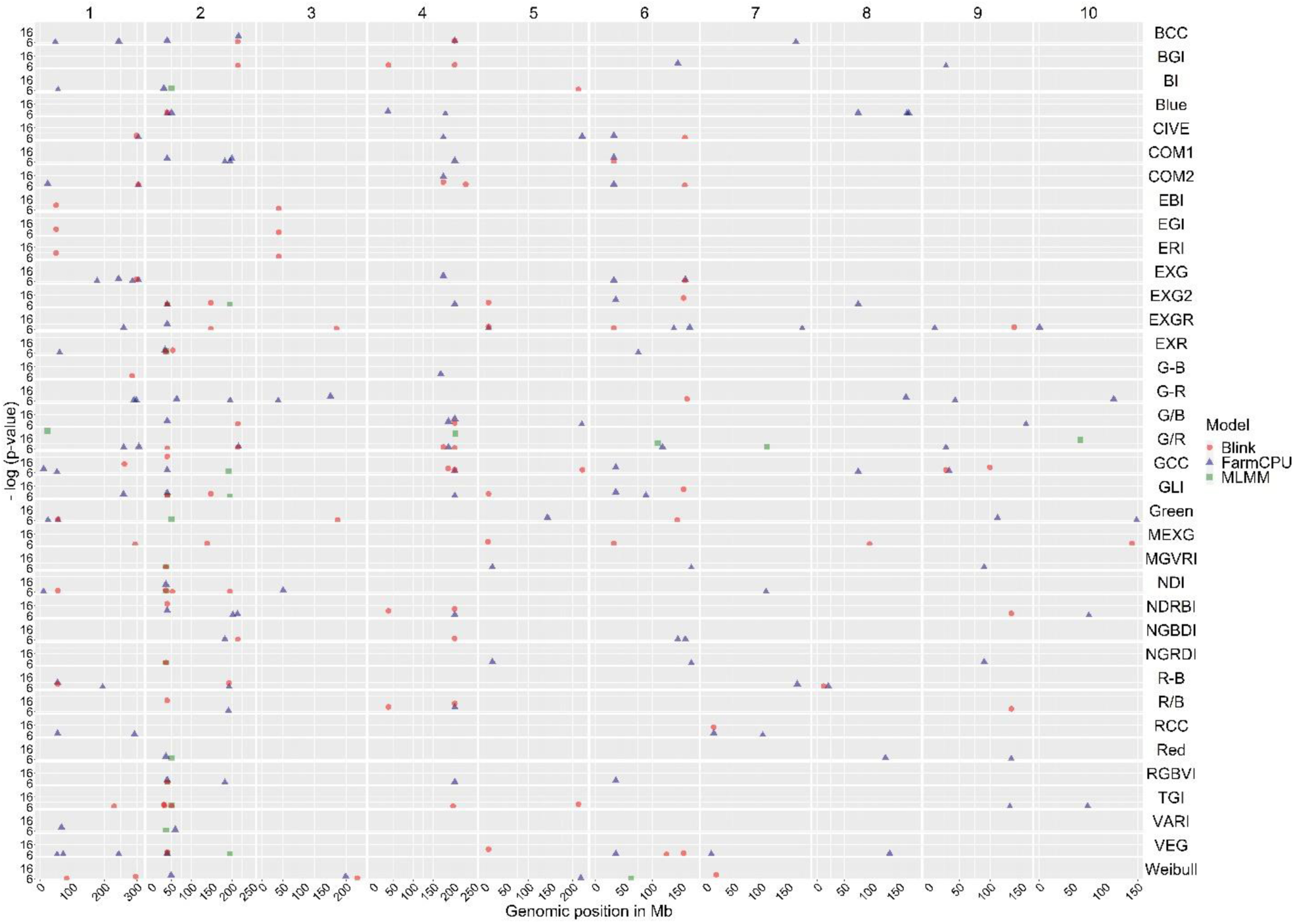
Combined Manhattan plots for each VI and Weibull _CHM (based on the right y-axis) in TPP_RGB. Below x-axis shows the genomic position of each chromosome (based on the above x-axis). Left y-axis shows the probability (-log10) of GWAS peaks between 6 to 26. Round, triangle and squares represent the Blink, FarmCPU and MLMM models results respectively.

**Figure S14.**
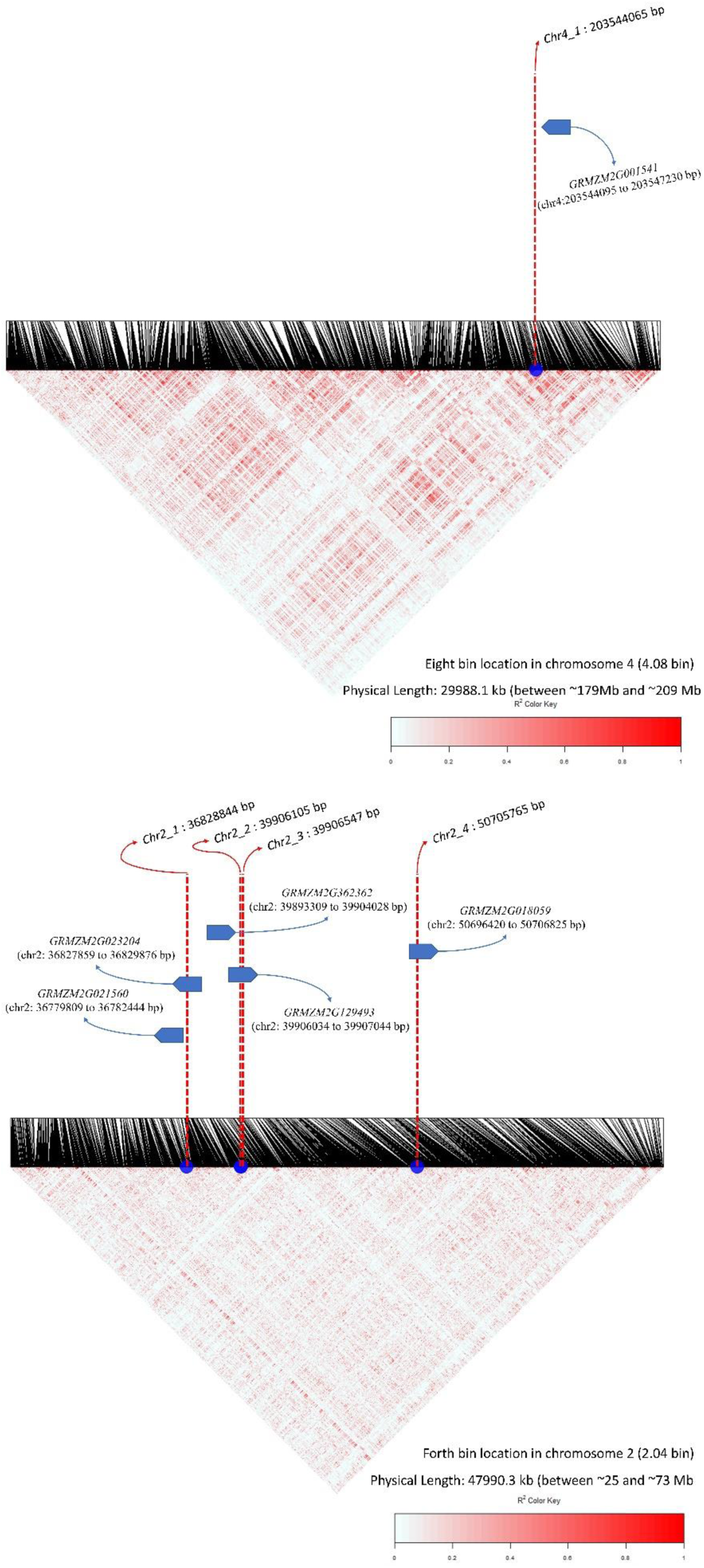
LD blocks around Chromosome two loci, *chr2_1, chr2_2, chr2_3, chr2_4* (below) and *chr4_1* (above) loci, nearby LD blocks of these loci within the 2.04 and 4.08 genomic bin locations and candidate gene annotations.

## References

1. J. L. Araus, S. C. Kefauver, M. Zaman-Allah, M. S. Olsen, J. E. Cairns, Translating high-throughput phenotyping into genetic gain. Trends in plant science 23, 451–466 (2018).

2. H. M. Lane, S. C. Murray, High Throughput can produce better decisions than high accuracy when phenotyping plant populations. Crop Science.

3. Y. Shi et al., Unmanned aerial vehicles for high-throughput phenotyping and agronomic research. PloS one 11, e0159781 (2016).

4. R. Bernardo, Predictive breeding in maize during the last 90 years. Crop Science (2021).

5. C. R. Henderson, Best linear unbiased estimation and prediction under a selection model. Biometrics, 423–447 (1975).

6. R. Bernardo, Prediction of maize single-cross performance using RFLPs and information from related hybrids. Crop Science 34, 20–25 (1994).

7. R. Bernardo, Best linear unbiased prediction of maize single-cross performance. Crop Science 36, 50–56 (1996).

8. R. Bernardo, Best linear unbiased prediction of the performance of crosses between untested maize inbreds. Crop Science 36, 872–876 (1996).

9. J. B. Endelman, Ridge regression and other kernels for genomic selection with R package rrBLUP. The plant genome 4 (2011).

10. T. H. Meuwissen, B. J. Hayes, M. E. Goddard, Prediction of total genetic value using genome-wide dense marker maps. Genetics 157, 1819–1829 (2001).

11. J. C. Whittaker, R. Thompson, M. C. Denham, Marker-assisted selection using ridge regression. Genetics Research 75, 249–252 (2000).

12. R. Bernardo, J. Yu, Prospects for genomewide selection for quantitative traits in maize. Crop Science 47, 1082–1090 (2007).

13. A. Adak et al., Validation of Functional Polymorphisms Affecting Maize Plant Height by Unoccupied Aerial Systems (UAS) Discovers Novel Temporal Phenotypes. G3 Genes| Genomes| Genetics (2021).

14. A. Adak et al., Unoccupied aerial systems discovered overlooked loci capturing the variation of entire growing period in maize. The Plant Genome, e20102 (2021).

15. A. Adak et al., Temporal Vegetation Indices and Plant Height from Remotely Sensed Imagery Can Predict Grain Yield and Flowering Time Breeding Value in Maize via Machine Learning Regression. Remote Sensing 13, 2141 (2021).

16. S. L. Anderson et al., Prediction of maize grain yield before maturity using improved temporal height estimates of unmanned aerial systems. The Plant Phenome Journal 2, 1–15 (2019).

17. J. A. Bac-Molenaar, D. Vreugdenhil, C. Granier, J. J. Keurentjes, Genome-wide association mapping of growth dynamics detects time-specific and general quantitative trait loci. Journal of experimental botany 66, 5567–5580 (2015).

18. M. T. Campbell et al., A comprehensive image-based phenomic analysis reveals the complex genetic architecture of shoot growth dynamics in rice (Oryza sativa). (2017).

19. M. J. Feldman et al., Time dependent genetic analysis links field and controlled environment phenotypes in the model C4 grass Setaria. PLoS genetics 13, e1006841 (2017).

20. B. Ward et al., High-throughput 3D modelling to dissect the genetic control of leaf elongation in barley (Hordeum vulgare). The Plant Journal 98, 555–570 (2019).

21. R. Wu, Z. Wang, W. Zhao, J. M. Cheverud, A mechanistic model for genetic machinery of ontogenetic growth. Genetics 168, 2383–2394 (2004).

22. S. Murray et al., G2F Maize UAV Data, College Station, Texas 2017. CyVerse Data Commons. doi 10 (2019).

23. S. L. Anderson, S. C. M. II, R/UAStools:: plotshpcreate: Create multi-polygon shapefiles for extraction of research plot scale agriculture remote sensing data. Frontiers in plant science 11 (2020).

24. F. I. Matias, M. V. Caraza-Harter, J. B. Endelman, FIELDimageR: an R package to analyze orthomosaic images from agricultural field trials. The Plant Phenome Journal 3, e20005 (2020).

25. X. Li, T. Guo, Q. Mu, X. Li, J. Yu, Genomic and environmental determinants and their interplay underlying phenotypic plasticity. Proceedings of the National Academy of Sciences 115, 6679–6684 (2018).

26. X. Liu, M. Huang, B. Fan, E. S. Buckler, Z. Zhang, Iterative usage of fixed and random effect models for powerful and efficient genome-wide association studies. PLoS genetics 12, e1005767 (2016).

27. V. Segura et al., An efficient multi-locus mixed-model approach for genome-wide association studies in structured populations. Nature genetics 44, 825–830 (2012).

28. M. Huang, X. Liu, Y. Zhou, R. M. Summers, Z. Zhang, BLINK: a package for the next level of genome-wide association studies with both individuals and markers in the millions. GigaScience 8, giy154 (2019).

29. O. N. Danilevskaya, X. Meng, Z. Hou, E. V. Ananiev, C. R. Simmons, A genomic and expression compendium of the expanded PEBP gene family from maize. Plant physiology 146, 250–264 (2008).

30. X.-h. Song et al., Integrating transcriptomic and proteomic analyses of photoperiod-sensitive in near isogenic maize line under long-day conditions. Journal of Integrative Agriculture 18, 1211–1221 (2019).

31. F. M. Aguate et al., Use of hyperspectral image data outperforms vegetation indices in prediction of maize yield. (2017).

32. R. J. Galán et al., Early prediction of biomass in hybrid rye based on hyperspectral data surpasses genomic predictability in less-related breeding material. Theoretical and Applied Genetics 134, 1409–1422 (2021).

33. M. R. Krause et al., Aerial high-throughput phenotyping enabling indirect selection for grain yield at the early-generation seed-limited stages in breeding programs. Crop Science, 3096–3114 (2020).

34. O. A. Montesinos-López et al., Predicting grain yield using canopy hyperspectral reflectance in wheat breeding data. Plant methods 13, 1–23 (2017).

35. J. Rutkoski et al., Canopy temperature and vegetation indices from high-throughput phenotyping improve accuracy of pedigree and genomic selection for grain yield in wheat. G3: Genes, Genomes, Genetics 6, 2799–2808 (2016).

36. J. Sun et al., High-throughput phenotyping platforms enhance genomic selection for wheat grain yield across populations and cycles in early stage. Theoretical and Applied Genetics 132, 1705–1720 (2019).

37. G. Wu, N. D. Miller, N. De Leon, S. M. Kaeppler, E. P. Spalding, Predicting Zea mays flowering time, yield, and kernel dimensions by analyzing aerial images. Frontiers in plant science 10, 1251 (2019).

38. C. D. Messina et al., Leveraging biological insight and environmental variation to improve phenotypic prediction: Integrating crop growth models (CGM) with whole genome prediction (WGP). European Journal of Agronomy 100, 151–162 (2018).

39. K. Kismiantini, O. A. Montesinos-López, J. Crossa, E. P. Setiawan, D. U. Wutsqa, Prediction of count phenotypes using high-resolution images and genomic data. G3 Genes| Genomes| Genetics (2021).

40. R. Rincent et al., Phenomic selection is a low-cost and high-throughput method based on indirect predictions: proof of concept on wheat and poplar. G3: Genes, Genomes, Genetics 8, 3961–3972 (2018).

## SI References

1. R. Escadafal, Remote sensing of soil color: principles and applications. Remote Sensing Reviews 7, 261–279 (1993).

2. K. Kraus, N. Pfeifer, Determination of terrain models in wooded areas with airborne laser scanner data. ISPRS Journal of Photogrammetry and remote Sensing 53, 193–203 (1998).

3. B. A. McFarland et al., Maize genomes to fields (G2F): 2014–2017 field seasons: genotype, phenotype, climatic, soil, and inbred ear image datasets. BMC research notes 13, 1–6 (2020).

4. J. C. Glaubitz et al., TASSEL-GBS: a high capacity genotyping by sequencing analysis pipeline. PloS one 9, e90346 (2014).

5. R. J. Elshire et al., A robust, simple genotyping-by-sequencing (GBS) approach for high diversity species. PloS one 6, e19379 (2011).

6. P. J. Bradbury et al., TASSEL: software for association mapping of complex traits in diverse samples. Bioinformatics 23, 2633–2635 (2007).

7. D. Money et al., LinkImpute: fast and accurate genotype imputation for nonmodel organisms. G3: Genes, Genomes, Genetics 5, 2383–2390 (2015).

8. J.-H. Shin, S. Blay, B. McNeney, J. Graham, LDheatmap: an R function for graphical display of pairwise linkage disequilibria between single nucleotide polymorphisms. Journal of statistical software 16, 1–10 (2006).

10. D. M. Woebbecke, G. E. Meyer, K. Von Bargen, D. A. Mortensen, Color indices for weed identification under various soil, residue, and lighting conditions. Transactions of the ASAE 38, 259–269 (1995).

11. P. J. Zarco-Tejada et al., Assessing vineyard condition with hyperspectral indices: Leaf and canopy reflectance simulation in a row-structured discontinuous canopy. Remote Sensing of Environment 99, 271–287 (2005).

12. A. J. Richardson, C. Wiegand, Distinguishing vegetation from soil background information. Photogrammetric engineering and remote sensing 43, 1541–1552 (1977).

13. T. Kataoka, T. Kaneko, H. Okamoto, S. Hata (2003) Crop growth estimation system using machine vision. in Proceedings 2003 IEEE/ASME International Conference on Advanced Intelligent Mechatronics (AIM 2003) (IEEE), pp b1079–b1083 vol. 1072.

14. M. Guijarro et al., Automatic segmentation of relevant textures in agricultural images. Computers and Electronics in Agriculture 75, 75–83 (2011).

15. J. M. Guerrero, G. Pajares, M. Montalvo, J. Romeo, M. Guijarro, Support vector machines for crop/weeds identification in maize fields. Expert Systems with Applications 39, 11149–11155 (2012).

16. M. R. Golzarian, R. A. Frick, Classification of images of wheat, ryegrass and brome grass species at early growth stages using principal component analysis. Plant Methods 7, 1–11 (2011).

17. G. E. Meyer, J. C. Neto, Verification of color vegetation indices for automated crop imaging applications. Computers and electronics in agriculture 63, 282–293 (2008).

18. G. Meyer, T. Hindman, K. Laksmi, MG (ed.), Deshazer JA, Machine vision detection parameters for plant species identification. Precision agri540 culture and biological quality, Boston, Massachusetts, USA 3, 3543 (1998).

19. M. Louhaichi, M. M. Borman, D. E. Johnson, Spatially located platform and aerial photography for documentation of grazing impacts on wheat. Geocarto International 16, 65–70 (2001).

20. X. P. Burgos-Artizzu, A. Ribeiro, M. Guijarro, G. Pajares, Real-time image processing for crop/weed discrimination in maize fields. Computers and Electronics in Agriculture 75, 337–346 (2011).

21. J. Bendig et al., Combining UAV-based plant height from crop surface models, visible, and near infrared vegetation indices for biomass monitoring in barley. International Journal of Applied Earth Observation and Geoinformation 39, 79–87 (2015).

22. E. R. Hunt, M. Cavigelli, C. S. Daughtry, J. E. Mcmurtrey, C. L. Walthall, Evaluation of digital photography from model aircraft for remote sensing of crop biomass and nitrogen status. Precision Agriculture 6, 359–378 (2005).

23. C. J. Tucker, Red and photographic infrared linear combinations for monitoring vegetation. Remote sensing of Environment 8, 127–150 (1979).

24. E. R. Hunt, C. Daughtry, J. U. Eitel, D. S. Long, Remote sensing leaf chlorophyll content using a visible band index. (2011).

25. A. A. Gitelson, Y. J. Kaufman, R. Stark, D. Rundquist, Novel algorithms for remote estimation of vegetation fraction. Remote sensing of Environment 80, 76–87 (2002).

26. T. Hague, N. Tillett, H. Wheeler, Automated crop and weed monitoring in widely spaced cereals. Precision Agriculture 7, 21–32 (2006).

27. C. S. Daughtry, C. Walthall, M. Kim, E. B. De Colstoun, J. McMurtrey Iii, Estimating corn leaf chlorophyll concentration from leaf and canopy reflectance. Remote sensing of Environment 74, 229–239 (2000).

28. D. Haboudane, J. R. Miller, E. Pattey, P. J. Zarco-Tejada, I. B. Strachan, Hyperspectral vegetation indices and novel algorithms for predicting green LAI of crop canopies: Modeling and validation in the context of precision agriculture. Remote sensing of environment 90, 337–352 (2004).

29. A. A. Gitelson, A. Vina, V. Ciganda, D. C. Rundquist, T. J. Arkebauer, Remote estimation of canopy chlorophyll content in crops. Geophysical Research Letters 32 (2005).

30. M. Vincini, E. Frazzi, P. D’Alessio, A broad-band leaf chlorophyll vegetation index at the canopy scale. Precision Agriculture 9, 303–319 (2008).

31. A. Huete et al., Overview of the radiometric and biophysical performance of the MODIS vegetation indices. Remote sensing of environment 83, 195–213 (2002).

32. R. E. Crippen, Calculating the vegetation index faster. Remote sensing of Environment 34, 71–73 (1990).

33. A. A. Gitelson, Y. J. Kaufman, M. N. Merzlyak, Use of a green channel in remote sensing of global vegetation from EOS-MODIS. Remote sensing of Environment 58, 289–298 (1996).

34. G. Rondeaux, M. Steven, F. Baret, Optimization of soil-adjusted vegetation indices. Remote sensing of environment 55, 95–107 (1996).

35. J.-L. Roujean, F.-M. Breon, Estimating PAR absorbed by vegetation from bidirectional reflectance measurements. Remote sensing of Environment 51, 375–384 (1995).

36. C. Buschmann, E. Nagel, In vivo spectroscopy and internal optics of leaves as basis for remote sensing of vegetation. International Journal of Remote Sensing 14, 711–722 (1993).

37. R. P. Sripada, R. W. Heiniger, J. G. White, A. D. Meijer, Aerial color infrared photography for determining early in-season nitrogen requirements in corn. Agronomy Journal 98, 968–977 (2006).

38. A. A. Gitelson, Wide dynamic range vegetation index for remote quantification of biophysical characteristics of vegetation. Journal of plant physiology 161, 165–173 (2004).

39. G. Le Maire, C. Francois, E. Dufrene, Towards universal broad leaf chlorophyll indices using PROSPECT simulated database and hyperspectral reflectance measurements. Remote sensing of environment 89, 1–28 (2004).

40. J. Qi, A. Chehbouni, A. R. Huete, Y. H. Kerr, S. Sorooshian, A modified soil adjusted vegetation index. Remote sensing of environment 48, 119–126 (1994).

41. B. Datt, Visible/near infrared reflectance and chlorophyll content in Eucalyptus leaves. International Journal of Remote Sensing 20, 2741–2759 (1999).

42. W. Wang et al., Estimating leaf nitrogen concentration with three-band vegetation indices in rice and wheat. Field Crops Research 129, 90–98 (2012).

43. J. M. Chen, Evaluation of vegetation indices and a modified simple ratio for boreal applications. Canadian Journal of Remote Sensing 22, 229–242 (1996).

44. D. Haboudane, J. R. Miller, N. Tremblay, P. J. Zarco-Tejada, L. Dextraze, Integrated narrow-band vegetation indices for prediction of crop chlorophyll content for application to precision agriculture. Remote sensing of environment 81, 416–426 (2002).

45. E. Barnes et al. (2000) Coincident detection of crop water stress, nitrogen status and canopy density using ground based multispectral data. in Proceedings of the Fifth International Conference on Precision Agriculture, Bloomington, MN, USA.

46. M. N. Merzlyak, A. A. Gitelson, O. B. Chivkunova, V. Y. Rakitin, Non-destructive optical detection of pigment changes during leaf senescence and fruit ripening. Physiologia plantarum 106, 135–141 (1999).

47. Q. Cao et al., Non-destructive estimation of rice plant nitrogen status with Crop Circle multispectral active canopy sensor. Field Crops Research 154, 133–144 (2013).

48. N. H. Broge, E. Leblanc, Comparing prediction power and stability of broadband and hyperspectral vegetation indices for estimation of green leaf area index and canopy chlorophyll density. Remote sensing of environment 76, 156–172 (2001).

49. C. F. Jordan, Derivation of leaf-area index from quality of light on the forest floor. Ecology 50, 663–666 (1969).

50. A. R. Huete, A soil-adjusted vegetation index (SAVI). Remote sensing of environment 25, 295–309 (1988).

51. J. Jasper, S. Reusch, A. Link, Active sensing of the N status of wheat using optimized wavelength combination: impact of seed rate, variety and growth stage. Precision agriculture 9, 23–30 (2009).

52. L. Sandham, H. Zietsman, Surface temperature measurement from space: a case study in the south western cape of South Africa. South African Journal of Enology and Viticulture 18, 25–30 (1997).

53. P. Gong, R. Pu, G. S. Biging, M. R. Larrieu, Estimation of forest leaf area index using vegetation indices derived from Hyperion hyperspectral data. IEEE transactions on geoscience and remote sensing 41, 1355–1362 (2003).

54. K. Erdle, B. Mistele, U. Schmidhalter, Comparison of active and passive spectral sensors in discriminating biomass parameters and nitrogen status in wheat cultivars. Field Crops Research 124, 74–84 (2011).

55. S. Elsayed, P. Rischbeck, U. Schmidhalter, Comparing the performance of active and passive reflectance sensors to assess the normalized relative canopy temperature and grain yield of drought-stressed barley cultivars. Field Crops Research 177, 148–160 (2015).

56. S. Ferrari, D. Vairo, F. M. Ausubel, F. Cervone, G. De Lorenzo, Tandemly duplicated Arabidopsis genes that encode polygalacturonase-inhibiting proteins are regulated coordinately by different signal transduction pathways in response to fungal infection. The Plant Cell 15, 93–106 (2003).

57. Z. Minic, Physiological roles of plant glycoside hydrolases. Planta 227, 723–740 (2008).

58. X. Wang et al., Genetic variation in ZmVPP1 contributes to drought tolerance in maize seedlings. Nature genetics 48, 1233–1241 (2016).

59. C. He et al., Early drought-responsive genes are variable and relevant to drought tolerance. G3: Genes, Genomes, Genetics 10, 1657–1670 (2020).

60. Z. Zhou et al., A QTL atlas for grain yield and its component traits in maize (Zea mays). Plant Breeding 139, 562–574 (2020).

61. J. Hattan, H. Kanamoto, M. Takemura, A. Yokota, T. Kohchi, Molecular characterization of the cytoplasmic interacting protein of the receptor kinase IRK expressed in the inflorescence and root apices of Arabidopsis. Bioscience, biotechnology, and biochemistry 68, 2598–2606 (2004).

62. X. Wu et al., Joint-linkage mapping and GWAS reveal extensive genetic loci that regulate male inflorescence size in maize. Plant Biotechnology Journal 14, 1551–1562 (2016).

63. Y. Wang, Y. Wang, X. Wang, D. Deng, Integrated meta-QTL and genome-wide association study analyses reveal candidate genes for maize yield. Journal of Plant Growth Regulation 39, 229–238 (2020).

64. L. Liu et al., KRN4 controls quantitative variation in maize kernel row number. PLoS genetics 11, e1005670 (2015).

